# The food grade bacterium *Lactobacillus helveticus VEL12193* promotes autophagy by releasing membrane vesicles

**DOI:** 10.1101/2024.10.13.618067

**Authors:** Marie-Agnès Bringer, Jana Al Azzaz, Bénédicte Buteau, Lil Proukhnitzky, Amaury Aumeunier, Simon Manceau, Luis G. Bermúdez-Humarán, Chain F Florian, Catherine Daniel, Elise Jacquin, Niyazi Acar, Aurélie Rieu, Pierre Lapaquette

## Abstract

Autophagy-related processes, including canonical macroautophagy, are crucial for maintaining cellular homeostasis in eukaryotic organisms. Alterations or reduced activity of these processes have been strongly linked to a broad range of human diseases including inflammatory bowel disease (IBD) and age-related diseases such as age-related macular degeneration - a disease that affect the central area of the retina. In contrast, long-term autophagy stimulation appears to be safe and to extend lifespan in model organisms such as mice. Thus, enhancing autophagy represents a promising strategy for promoting healthy aging. Several studies indicate that the gut microbiota can influence host autophagy at the gut mucosa but also in peripheral organs, and some microbial metabolites have been identified as autophagy modulators. In this study, we studied the capacity of bacterial species commonly used in food fermentation (ferments) or health (probiotics) to modulate host autophagy by *in vitro* and *in vivo* approaches. *In vitro* screening of a library of 11 bacterial strains revealed a strain-dependent ability of lactobacilli and bifidobacteria to stimulate autophagy in human epithelial cells. The *Lactobacillus helveticus* strain VEL12193, isolated from cheese, emerged as the most effective inducer of autophagy. *In vivo* experiment using mice showed that long-term dietary supplementation with *L. helveticus* VEL12193 was associated with stimulation of autophagy in the gut mucosa and retina. We identified *L. helveticus*-derived membrane vesicles (MVs) as a bacterial component involved in bacterial-induced autophagy in epithelial and immune cells. Moreover, *in vitro*, we demonstrated that *L. helveticus* VEL12193 possesses immunomodulatory properties in macrophages, as well as in the gut mucosa of a preclinical mouse model of IBD. With this study we provide robust proof of concept that ferments/probiotics can stimulate autophagy at the organism scale and that this phenotype involved MVs. In addition, we identify *L. helveticus* VEL12193 as a candidate strain of interest for the design of healthy-aging strategies.

## Introduction

Autophagy is a ubiquitous highly dynamic eukaryotic cellular process mediating the lysosomal degradation of intracellular cargoes (e.g., organelles, proteins, lipids or intracellular pathogens). The activity level of this process depends on cell type and varies in response to a wide range of stresses, including metabolic and immune perturbations, to maintain cellular homeostasis. At molecular level, orchestration of the autophagy machinery relies on more than 40 autophagy-related (ATG) proteins with various activities (e.g., kinases, ubiquitin-like conjugation enzymes or lipid transfer proteins). Several types of autophagy can be distinguished. Macroautophagy (called autophagy hereafter) refers to the formation of a crescent-shaped membrane structure (called phagophore or isolation membrane) that elongates around the cargoes, and finally closes to form a double-membrane vacuole called the autophagosome [1]. Beyond canonical autophagy described above, a growing number of studies describes several non-canonical forms of autophagy involving only a subset of ATG proteins and fulfilling various roles in cells notably in cellular membrane dynamic [2,3].

Autophagy is a keystone of homeostasis by regulating various biological processes such as energy balance, response to oxidative stress, inflammation and the cell death/survival balance. A decline in autophagy has been repeatedly reported in various organs and model organisms during aging, and dysfunction of this homeostasis gatekeeper is thought to impact multiple features of aging and contributes to the development of age-related diseases such as age-related macular degeneration (AMD). AMD is a chronic eye disease affecting the macula - a small retinal area involved in central vision – which is the leading cause of visual impairment and severe vision loss in western countries with 288 million people affected worldwide by 2040 and for which the therapeutic arsenal is limited and non-curative [4–7]. In the retina, autophagy is a crucial process that sustains the function and homeostasis of both the neuroretina and the retinal pigmentary epithelium, which have a high metabolic activity and are particularly exposed to oxidative stress. Alterations of retinal autophagy have been identified in AMD patients and they are suspected to be a key contributor to the disease [8–11]. Besides age-associated diseases, defects in autophagy have also been associated with the onset of other human diseases, including inflammatory bowel diseases (IBDs) such as Crohn’s disease for which no curative treatment are available [12]. Dysfunction of autophagy in the gut mucosa – a phenotype that can result from polymorphisms and/or mutations in genes related to autophagy (e.g., *ATG16L1*, *IRGM*, *NDP52* or *ULK1*) in patients with Crohn’s disease - has been reported to be associated with uncontrolled innate immunity responses including defects in intracellular bacterial killing, decreased antimicrobial peptide secretion, impaired antigen presentation and exacerbated inflammatory response, which are factors contributing to the disease [13–16].

The predicted continued rise in the prevalence of IBD and age-related diseases such as AMD makes it crucial to find new levers and develop new strategies for preventing these diseases. We can assume that strategies focused on supporting autophagy could be beneficial in the context of diseases such as AMD and IBD that involve autophagy defects. Indeed, chronic and moderate stimulation of autophagy has been shown to increase lifespan and favor healthy aging in various model organisms, suggesting that long-term activation of autophagy is relatively safe and can provide health benefits [17,18]. However, dietary or therapeutic intervention specifically designed to modulate autophagy for health benefits that are safe are not available so far. Autophagy was described historically in yeast as a sensor of starvation and evidence indicates that intermittent fasting or calorie restriction in mammals induces autophagy and increase longevity [19]. However, it has been shown that such practice can have harmful effects at the cell level [20] but also if uncontrolled at the organism level (e.g., undernourishment, dehydration). Moreover, several patented pharmacological modulators of autophagy have been described. But, in all cases, their precise specificity is not known and many of them, such as rapamycin analogues or niclosamide, act on multiple intracellular pathways, increasing the risk of harmful effects [16,21]. Thus, developing innovative approaches stimulating this process remains essential to be able to propose new strategies to prevent autophagy decline and promote healthy aging.

Interestingly, the gut microbiota can influence autophagy both locally in the gut and remotely in peripheral organs, positioning gut microbes as a promising lever to boost autophagy at the scale of the organism [22]. Several microbial-derived products including pathogen-associated molecular patterns (PAMPs; e.g., peptidoglycan, lipoteichoic acids, etc.) have been shown to modulate autophagy *in vitro* and *in vivo via* their interaction with pathogen recognition receptors (PRRs; e.g., Toll-like receptors, Nod-like receptors, etc.) [22–24]. In addition, metabolites produced by microorganisms acting on host metabolism can also influence autophagy, as exemplified *in vivo* by the inhibitory effect on gut autophagy of the short chain fatty acid (SCFA) butyrate or the stimulatory effect on autophagy of lactate in skeletal muscles [25,26]. In addition to bacteria, fungi can also exhibit pro-autophagic activities in host cells through cell wall compounds such as β-glucans or *via* the production of metabolites like the disaccharide trehalose [27,28]. However, although some microbial factors and their associated host cell receptors involved in autophagy modulation have been identified, characterization of microorganisms displaying pro-autophagic activities still requires functional screening *in vitro* and *in vivo* validation. Indeed, the regulation of autophagy is cell type dependent and complex as it involves multiple signaling pathways.

In this study, we analyzed the potential of 11 food-grade bacterial strains belonging to bacterial species frequently used as probiotics - lactobacilli and bifidobacteria - to stimulate autophagy. We first screened this bacterial library *in vitro* using human epithelial cells and analyzing several markers of autophagy activation by different approaches. This robust screening led us to select one strain - the *Lactobacillus helveticus* strain VEL12193 - with a promising potential for inducing autophagy. Next step was to validate the pro-autophagic potential of this bacterial strain in an *in vivo* model and in two locations, the gut mucosa and the retina, the latter constituting a proof of concept of a distant effect from the gut. We showed that long-term consumption (6 months) of a diet supplemented with *L. helveticus* VEL12193 resulted in autophagy activation both in the gut mucosa and in the retina of old mice (18-month-old at the end of the experiment). Furthermore, we identified membrane vesicles (MVs) released by *L. helveticus* VEL12193 as a bacterial-derived product that stimulates autophagy *in vitro* in human epithelial and immune cells. In addition to their pro-autophagic activity, we also showed that *L. helveticus*-derived MVs had immunomodulatory properties *in vitro* by decreasing the expression of pro-inflammatory cytokines in lipopolysaccharide-stimulated cells. This immunomodulatory effect of *L. helveticus* has been validated *in vivo* in dextran sulfate sodium (DSS)-treated mice, a model of colitis.

## Results

### In vitro screening of lactobacilli and bifidobacteria strains to stimulate autophagy

Eleven Gram positive bacteria belonging to the lactobacilli or *Bidifobacterium* families were analyzed for their ability to modulate autophagy in human epithelial cells (**Table 1**). To increase the robustness of this phenotypic analysis, we used several approaches. First, we compared the level of autophagy activation induced by the different bacterial strains by counting the number of intracellular vacuoles (appearing as dots on images) positive for two autophagy markers - LC3 and WIPI2 - in HeLa GFP-LC3 reporter cells by imaging. The LC3 proteins are described to be present on autophagic vacuoles from initial to late stages of the process whereas WIPI2 proteins are only present on the isolation membrane, at initial stages [29]. We observed that two strains, *Bifidobacterium longum* LBH422 and *L. helveticus* VEL12193, induced a significant increase in both the number of LC3 and WIPI2 dots compared to untreated cells (**Figure 1A** and **1B**). *B. longum* BH422 and *L. helveticus* VEL12193 were also associated with a significant increase in the number of WIPI2-positive vacuoles, compared to untreated cells (**Figure 1A** and **1C**). As illustrated with *L. helveticus* VEL12193 strain-treated cells, co-localization of the two markers were sometimes observed indicating the presence of nascent autophagosomes (**Sup Figure 1**). Of note, none of the tested strains decreased the number of LC3 or WIPI2 dots per cell compared with untreated cells, suggesting that these bacteria have no inhibitory effects on autophagy, at least at initial steps of the process.

**Figure 1.**
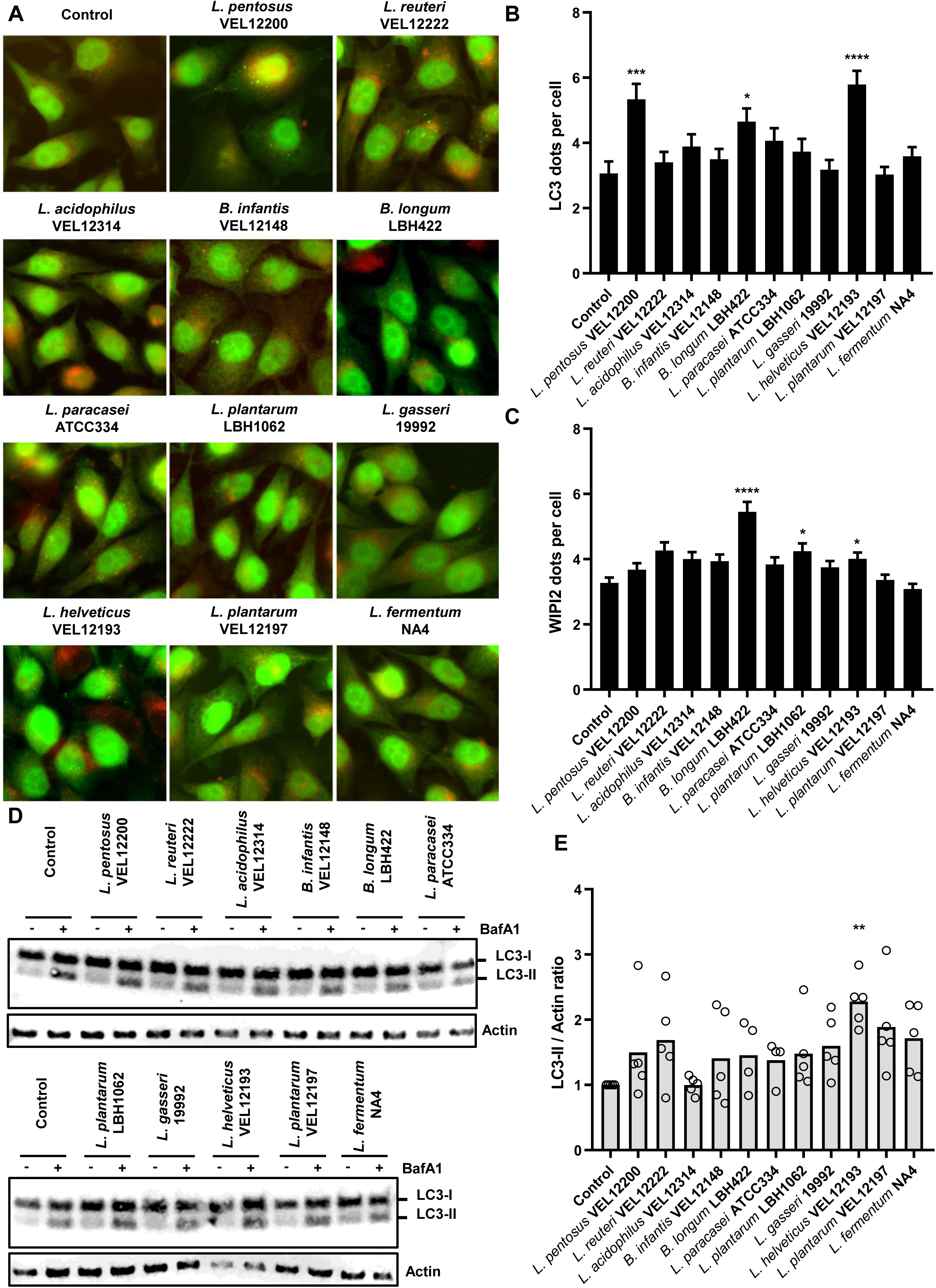
Strain-dependent stimulation of autophagy in Hela GFP-LC3 cells. **(A)** Representative images of GFP-LC3 HeLa cells treated or not (control) for 2 h with 11 different strains of lactobacilli and bifidobacteria. Cells were immunostained with anti-WIPI2 antibodies (red). LC3 protein coupled to GFP (GFP-LC3) appears in green. (**B** and **C**) Quantification of the number of (**B**) LC3 and (**C**) WIPI2 dots per cell 2 h after treatment of GFP-LC3 Hela cells with bacteria. Data are mean +/- SEM of three independent experiments. (**D** and **E**) *In vitro* autophagy flux assay. (**D**) Representative immunoblots of GFP-LC3 HeLa cells treated or not (control) with 11 different strains of lactobacilli and bifidobacteria for 2 h, in presence or absence of the autophagy flux inhibitor bafilomycin A1 (BafA1). Immunoblot analyses were performed using anti-LC3B and anti-actin antibodies. (**E**) The LC3-II/Actin ratio was calculated and normalized to that obtained in control cells without BafA1 (n=5). (**B**, **C** and **E**) Kruskal-Wallis test (* *p*<0.05, *** *p*<0.001 and **** *p*<0.0001).

**Table 1.**
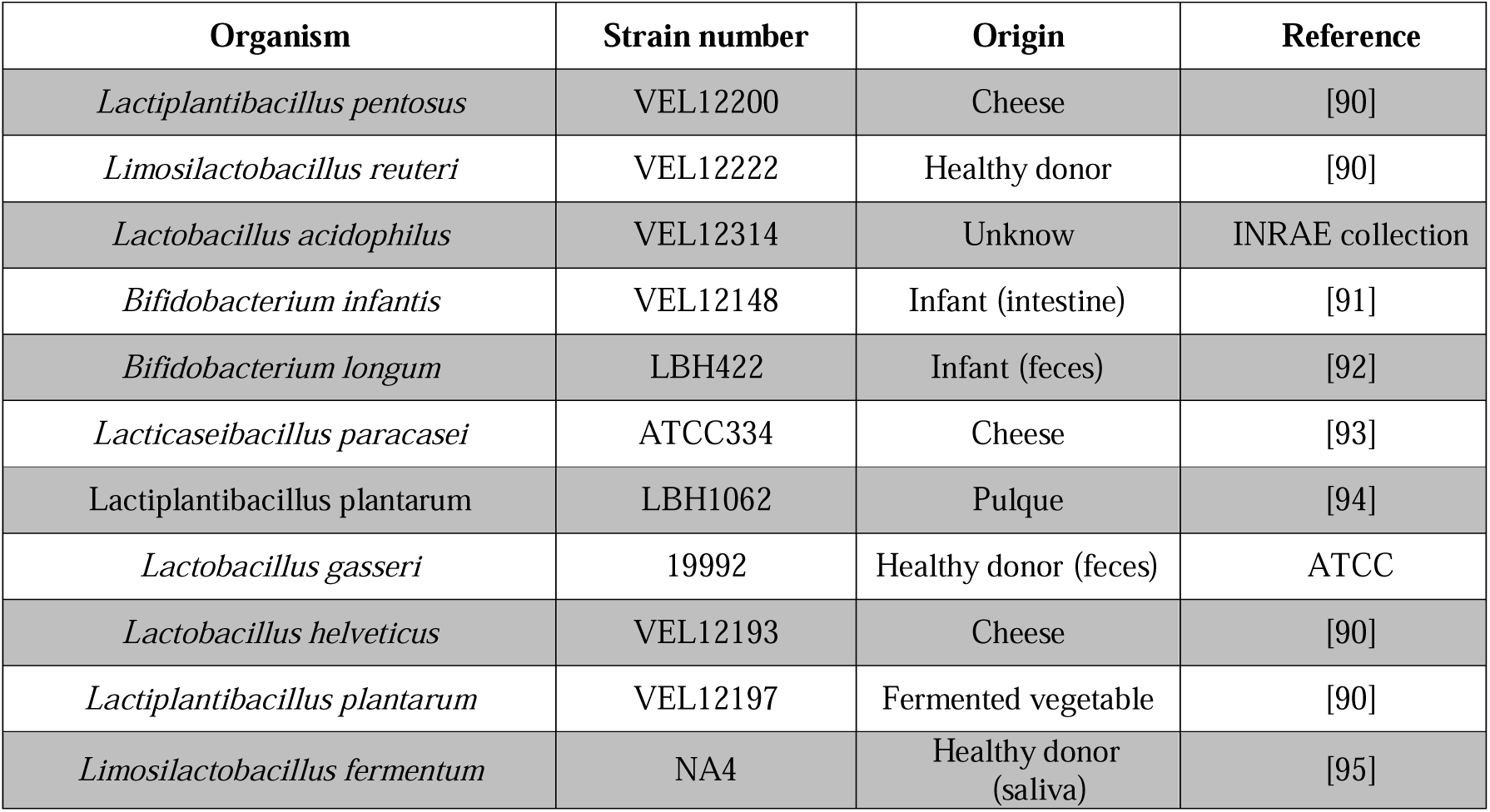
Bacterial strains used in this study.

The marked increase in LC3 dots associated with *B. longum* LBH422 and *L. helveticus* VEL12193 cell treatment could be due either to an induction of the autophagy process (i.e., formation of new autophagosomes) or a blockage in the late step of the process (inhibition of autophagosomes maturation), resulting in accumulation of autophagy vacuoles and thus in LC3. To avoid misinterpretation, we next analyzed the autophagic flux using bafilomycin A1 (BafA1), an autophagy flux inhibitor that prevents autophagosome-lysosome fusion and assessing the accumulation rate of endogenous LC3-II - the lipidated form of LC3 that is recruited on autophagy vacuole - by western blot [30]. Accumulation of LC3-II in cells treated with bacterial strains compared to untreated cells would suggest a stimulation of the autophagy process. We observed that only treatment with *L. helveticus* VEL12193 strain led to a significant accumulation of LC3-II compared to untreated HeLa cells (**Figure 1D** and **1E**).

Altogether, these results show that not all strains of lactobacilli and bifidobacteria have the capacity to stimulate autophagy in epithelial cells *in vitro*. Among the 11 tested strains, *L. helveticus* strain VEL12193 was identified by the various approaches used as being a promising autophagy inducer.

To validate the pro-autophagy property of *L. helveticus* VEL12193 and to make our results even more relevant, we repeated similar experiments in the human intestinal epithelial cell line HCT116. These cells are commonly used to study autophagy [3,31,32]. A significant increase in the number of intracellular vacuoles positive for LC3 and WIPI2 was observed in HCT116 cells treated with *L. helveticus* VEL12193 (**Figure 2A-C**). In addition, a significant increase in the LC3-II accumulation was observed in *L. helveticus* VEL12193- and BafA1-treated HCT116 cells compared to untreated cells (**Figure 2D** and **2E**). These results confirm the pro-autophagic property of *L. helveticus* VEL12193 *in vitro*.

**Figure 2.**
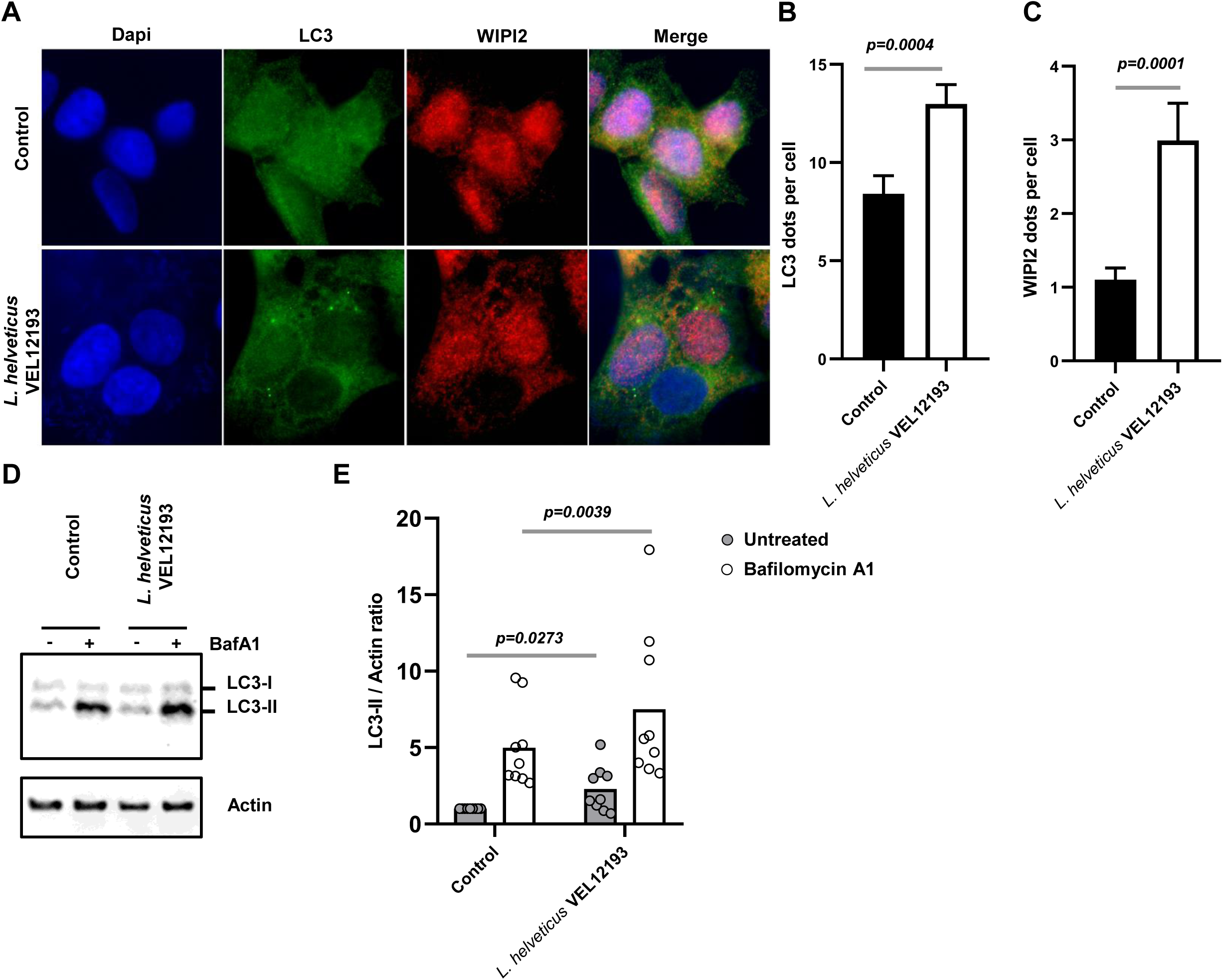
L. helveticus strain VEL12193 stimulates autophagy in HCT116 human intestinal epithelial cells. **(A**) Representative images of HCT116 cells treated or not (control) for 2 h with the *L. helveticus* strain VEL12193. Cells were immunostained with anti-WIPI2 (red), and anti-LC3 (green) antibodies. Nuclei were stained with DAPI (blue). (**B** and **C**) Quantification of the number of (**B**) LC3 and (**C**) WIPI2 dots per cell 2 h after treatment of HCT116 cells with bacteria. Data are mean +/- SEM of three independent experiments. Mann-Whitney test was used and *p*-values are indicated on the graphs. (**D** and **E**) *In vitro* autophagy flux assay. (**D**) Representative immunoblots of HCT116 treated or not (control) with *L. helveticus* strain VEL12193 for 2 h, in presence or absence of bafilomycin A1 (BafA1). Immunoblot analyses were performed using anti-LC3B and anti-actin antibodies. (**E**) The LC3-II/Actin ratio was calculated and normalized to that obtained in control cells without BafA1 (n=9). Wilcoxon matched pairs signed rank test was used, and *p*-values are indicated on the graphs.

### Effect of a long-term dietary supplementation of L. helveticus VEL12193 in mice on intestinal autophagy

Next, we explored whether *L. helveticus* strain VEL12193 can enhance autophagy *in vivo* in mice. As the intestinal compartment is the one through which bacteria-supplemented food will transit, we first focus on autophagy in the gut mucosa. Since activity of the autophagy process declined with aging [4], we evaluated the ability of a long-term supplementation (6 months) to stimulate autophagy in aged mice (18-month-old at the end of the experiment). For such long-term protocol, it was not conceivable ethically to administrate the bacteria by oral gavage. Thus, we chose to bring the bacteria along with the food by formulating pellets enriched with *L. helveticus* VEL12193. Supplementation of *L. helveticus* VEL12193 strain was well-tolerated by mice with no noticeable side effects, including feces consistency as previously reported [33]. Diet supplementation with *L. helveticus* was associated with a significant increase in the proportion of *L. helveticus* DNA found in the mice feces compared to mice fed control diet (**Sup Figure 2A**). No effect of *L. helveticus* VEL12193 supplementation on weight gain was observed (**Sup Figure 2B**).

Autophagy is a highly dynamic process whose activity remains difficult to assess *in vivo*. [34]. As the intestinal compartment is the one through which bacteria-supplemented food transit, we first studied autophagy in the mouse gut mucosa. Analysis of the autophagy flux in colonic explants revealed that long-term consumption of food enriched with *L. helveticus* VEL12193 significantly increased the level of LC3-II in the colon both at the basal level (without treatment with autophagy inhibitors, leupetine (Leu) and NH_4_Cl) and when the autophagy flux is blocked by inhibitors (Leu/ NH_4_Cl) compared to control mice (**Figure 3A** and **3B**). The accumulation trend of the p62 protein - a receptor of autophagy that is degraded by the process - in explants treated with autophagy flux inhibitors compared to untreated explant confirmed the efficacy of the autophagy flux blockade by the drugs. However, we observed no significant difference related to the dietary supplementation of mice with *L. helveticus* VEL12193 (**Figure 3A** and **3C**).

**Figure 3.**
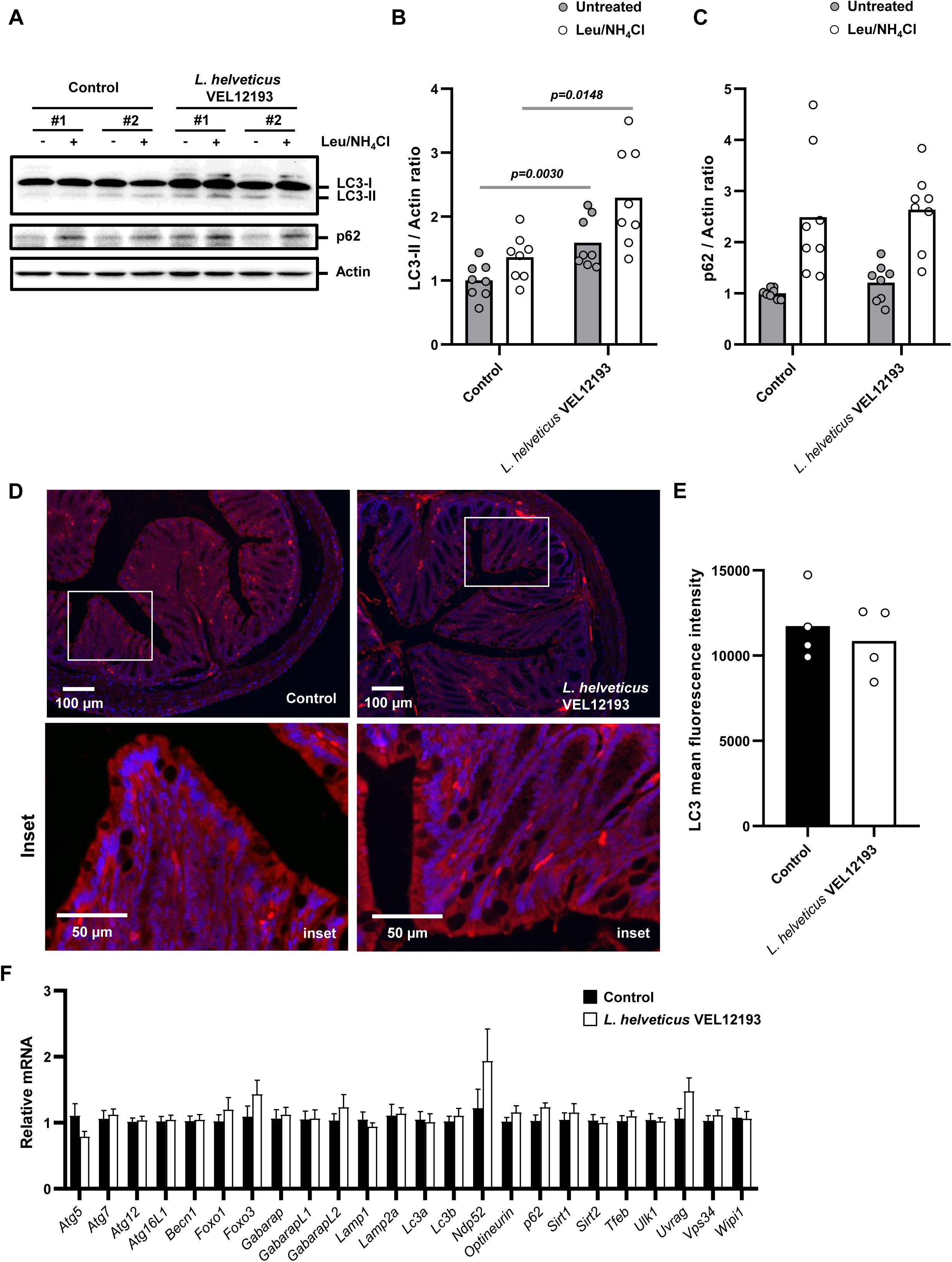
Long-term dietary supplementation of mice with L. helveticus VEL12193 prevents autophagy decline during aging in the gut mucosa. (**A-C**) *Ex vivo* autophagy flux assay. (**A**) Representative immunoblots of colonic explants from mice fed a control or *L. helveticus* VEL12193-supplemented diet incubated or not *ex vivo* with autophagy inhibitors (leupeptin (Leu) and NH_4_Cl). Immunoblot analyses were performed using anti-LC3B, anti-p62 and anti-actin antibodies. (**B** and **C**) The LC3-II/Actin (**B**) and p62/Actin ratio (**C**) were calculated and normalized to those obtained in control colonic explants untreated with autophagy flux inhibitors (n=8). Mann-Whitney test was used and *p*-values are indicated on the graphs. (**D**) Representative images of colon sections from mice fed a control or *L. helveticus* VEL12193-supplemented diet. White squares in upper panels indicate inset areas displayed in the corresponding lower panels. Samples were immunostained with anti-LC3 antibodies (red) and nuclei were stained with Hoescht (blue). (**E**) The mean intensity of LC3 staining in the gut mucosa sections was determined using ImageJ software. n=4 mice/group (Five fields per mouse section). Mann-Whitney test was used. (**F**) Expression of genes encoding enzymes involved in the autophagy pathway in the colon of mice fed a control diet (n=9) or fed a diet supplemented with *L. helveticus* VEL12193 (n=8). The levels of mRNA were normalized to *Hprt* mRNA level for calculation of the relative levels of transcripts. Data are mean +/- SEM. Mann-Whitney test was used.

It is technically difficult to assess subtle changes in the number of autophagy vacuoles by immunofluorescence on tissue sections. Immunostaining of colon sections with LC3 antibodies revealed a heterogeneous expression of this marker in the gut mucosa characterized by the presence of cells highly positive for this autophagy marker. Analysis of the global mean fluorescence intensity of LC3 did not show significant difference between colon sections of *L. helveticus* VEL12193 supplemented mice and those of untreated mice (**Figure 3D** and **3E**).

Since autophagy is regulated at the transcriptional level [35], we analyzed the expression levels of 24 genes encoding autophagy related proteins (ATGs) in the colonic mucosa of mice supplemented with *L. helveticus* and control mice (**Figure 3F**). A 6-month dietary supplementation with *L. helveticus* was not associated with modulation of the expression of ATG genes including genes encoding proteins involved in the transcriptional control of autophagy (*Foxo1*, *Foxo3*, *Sirtuin 1*, *Sirtuin 2* and *Tfeb*), in initiation steps (*Becn1*, *Ulk1*, *Uvrag*, *Vps34* and *Wipi1*), in selective forms of autophagy (*Ndp52*, *Optineurin* and *p62*), in conjugation systems required for elongation of the phagophore (*Atg5*, *Atg7*, *Atg12*, *Atg16l1*, *Gabarap*, *GabarapL1*, *GabarapL2*, *Lc3a* and *Lc3b*) and in maturation (*Lamp1* and *Lamp2a*).

These results did not show any transcriptional activation of autophagy associated with long-term consumption of *L. helveticus* VEL12193 but revealed a stimulation of the colonic autophagy flux in the colons of 18-month-old mice indicating that consumption of this bacterial strain supported autophagy activity in the gut mucosa of old mice.

### Impact of long-term consumption of *L. helveticus* VEL12193 on autophagy at the extraintestinal level: the retina as a study model and proof of concept

Evidence has accumulated these last decades showing that gut microbiota influence autophagy in peripheral organs [22,36]. Thus, we studied whether long-term dietary supplementation of mice with *L. helveticus* VEL12193 modulates autophagy in mouse retina (**Figure 4**). *Ex vivo* analysis of the autophagy flux in the retinas of 18-month-old mice supplemented or not with *L. helveticus* was analyzed. Significant increases in the basal level of LC3-II and a near significant accumulation of LC3-II in condition where the autophagy flux is blocked by autophagy inhibitors (Leu/ NH_4_Cl) were observed in retinas of *L. helveticus*-supplemented mice compared to those of control mice (**Figure 4A** and **4B**). Similarly to what we observed in the colonic mucosa, treating cells with autophagy inhibitors resulted in an increase in p62 level compared to untreated cells indicating that the autophagy flux was blocked by the drugs. However, in this organ again long-term consumption of *L. helveticus* VEL12193 had no effect on p62 accumulation (**Figure 4A** and **4C**).

**Figure 4.**
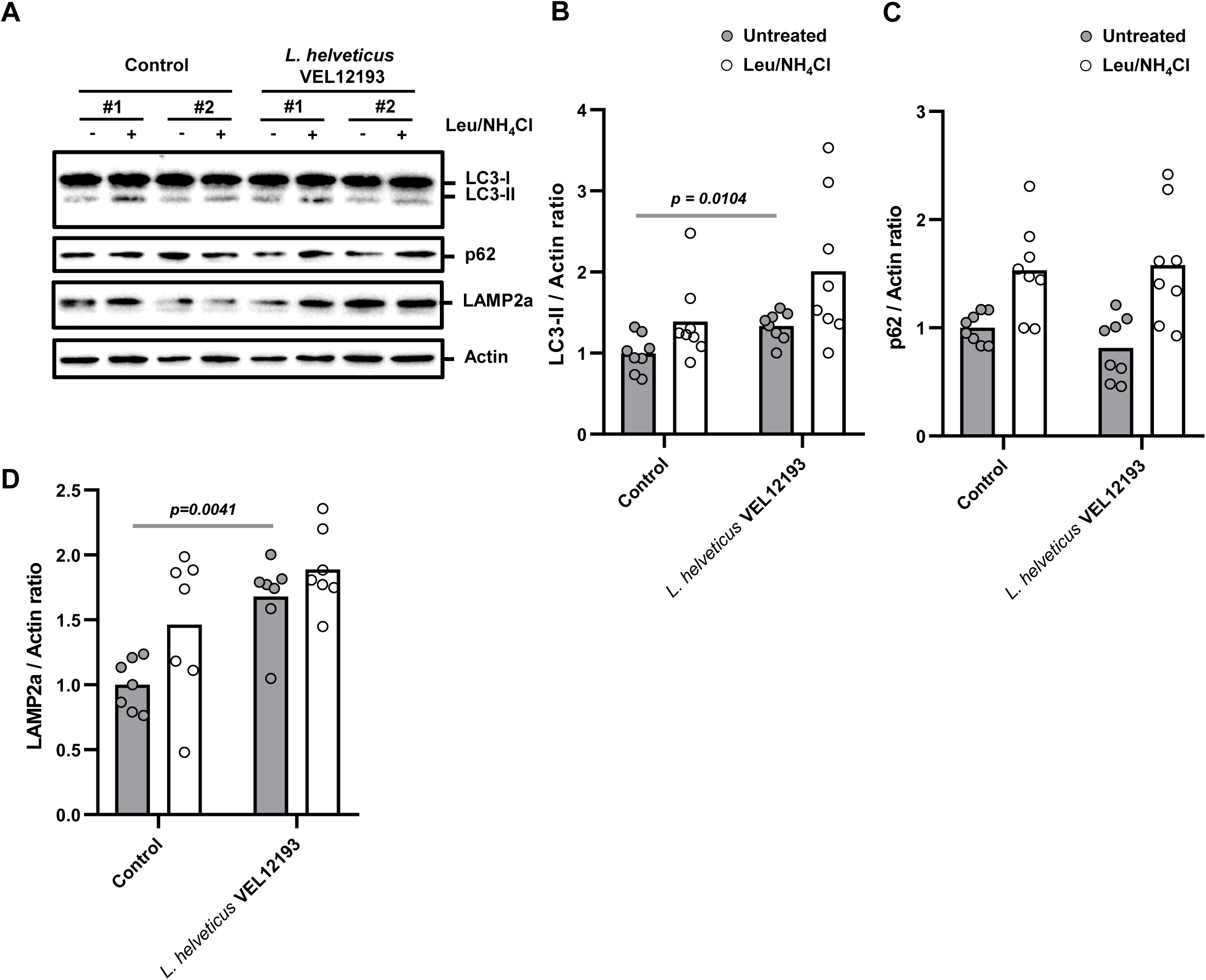
L. helveticus VEL12193 stimulates autophagy in the retina. (**A-D**) *Ex vivo* autophagy flux assay. Representative immunoblot of retinal explants from mice fed a control or *L. helveticus* VEL12193-supplemented diet incubated or not *ex vivo* with autophagy inhibitors (leupeptin (Leu) and NH_4_Cl) for 4 h. Immunoblot analyses were performed using anti-LC3B, anti-p62, anti-LAMP2a and anti-actin antibodies. (**B, C** and **D**) The LC3-II/actin (**B**) p62/actin (**C**) and LAMP2a/actin (**D**) ratio were calculated and normalized to those level obtained in control retinal explants untreated with autophagy flux inhibitors (n=8). Mann-Whitney test was used and *p*-values are indicated on the graphs.

Chaperone-mediated autophagy (CMA) is a form of autophagy that ensures the selective degradation of cellular proteins tagged with a KFERQ-like motif by lysosomes [37]. Crosstalk regulation exists between macroautophagy (generically referred to as autophagy) and CMA [38][38][38]. Interestingly, it has been reported that the reduction in macroautophagy activity observed during aging in the retina coincides with an increase in CMA. This cross talk is unidirectional in retina cells and is thought to preserve retinal homeostasis. The lysosomal protein LAMP2a is commonly used as a marker of CMA [39,40]. Interestingly, the basal level of LAMP2a was significantly higher in retinas of *L. helveticus*-supplemented mice, compared to those of control mice (**Figure 4A** and **4D**).

Altogether, these results suggest that long-term consumption of *L. helveticus* VEL12193 support both macroautophagy and CMA in the retina of old mice [41,42].

### Involvement of membrane vesicles (MVs) released by L. helveticus VEL12193 to stimulate autophagy

Health benefits associated with lactic acid bacteria are at least in part mediated by compounds or metabolites produced and secreted by these bacteria [43]. To gain insight into the molecular mechanisms by which *L. helveticus* VEL12193 stimulates autophagy, we have analyzed the ability of *L. helveticus* VEL12193 culture supernatant (SN) to induce such effect.

For this purpose, the cell-free supernatant of *L. helveticus* (SN *Lh*) and its growth medium (MRS) as a control were concentrated by size-exclusion ultrafiltration with a 3 kDa (> 3 kDa) or a 100 kDa (> 100 kDa) molecular weight cutoff. Treatment of GFP-LC3 Hela cells with the SN *Lh* > 3 kDa fraction was associated with a significant increase in the number of GFP-LC3 positive dots (**Figure 5A** and **5B**) and an increase trend upon Bafilomycin A1 treatment in the accumulation of LC3-II compared to cells treated with the control fraction (>3 kDa from MRS) (**Figure 5C** and **5D**). Interestingly, these phenotypes were still observed with the >100 kDa fraction (**Figure 5A-D**), suggesting that a bacterial compound with a high molecular weight was involved in the ability of *L. helveticus* to stimulate autophagy.

**Figure 5.**
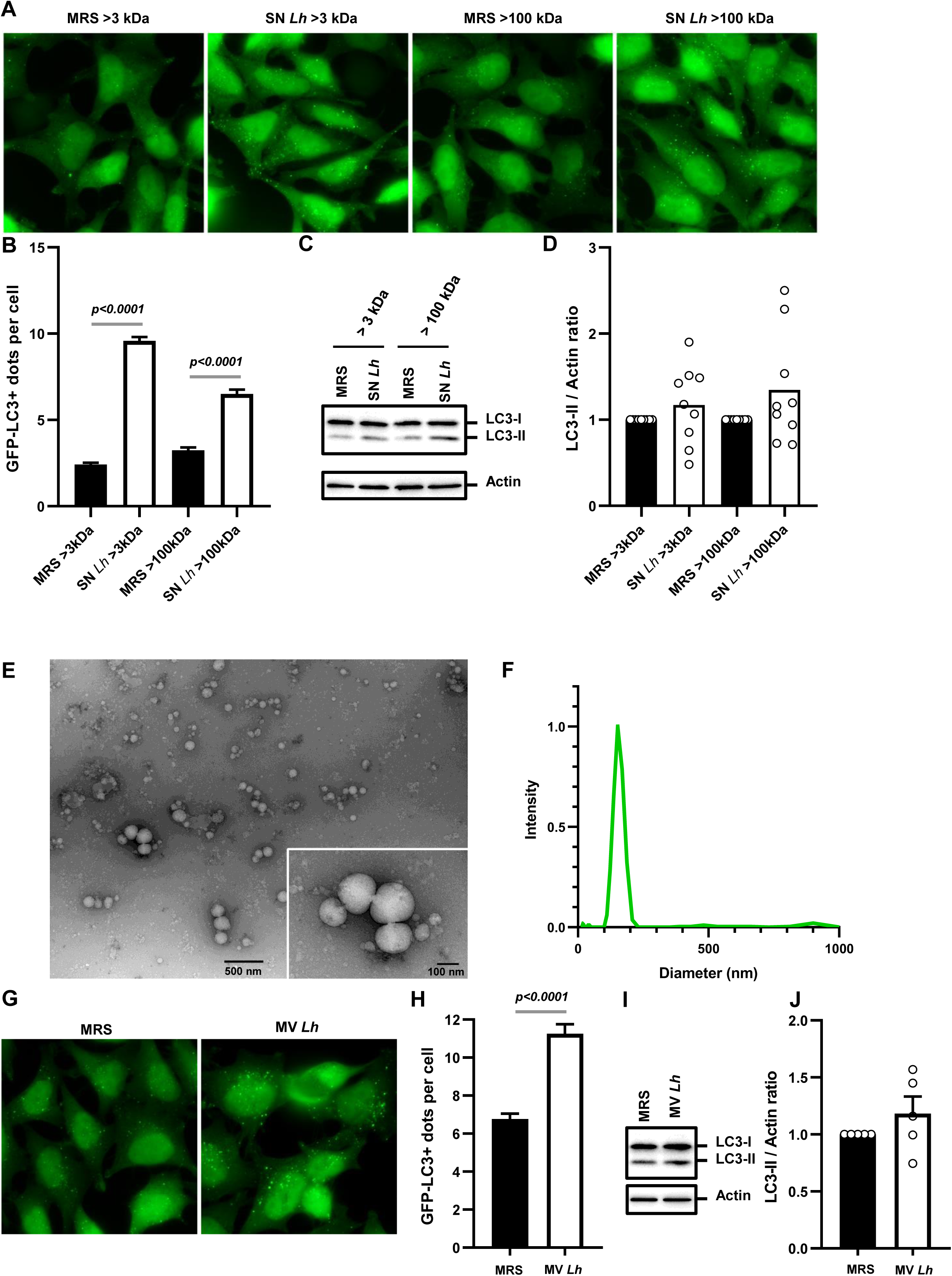
Membrane vesicles released by L. helveticus VEL12193 stimulate autophagy in epithelial cells. **(A)** Representative images of GFP-LC3 HeLa cells treated for 2 h with fractions (>3kDa or >100kDa) of *L. helveticus* supernatant (SN *Lh*) or MRS medium (bacterial culture medium,control). LC3 protein coupled to GFP (GFP-LC3) appears in green. (**B**) Quantification of the number of LC3 dots per cell at 2 h post treatment in GFP-LC3 Hela cells. Data are mean +/- SEM of three independent experiments. Mann-Whitney test was used and *p*-values are indicated on the graph. (**C**) *In vitro* autophagy flux assay. Representative immunoblots of GFP-LC3 HeLa cells treated or not (control) with the fractions (>3kDa or >100kDa) of *L. helveticus* SN (SN *Lh*) for 2 h, in presence of the autophagy inhibitor bafilomycin A1 (BafA1). Immunoblot analyses were performed using anti-LC3B and anti-actin antibodies. (**D**) The LC3-II/Actin ratio was calculated and normalized to that obtained in control cells without BafA1 (n=9). Wilcoxon matched pairs signed rank test was used. (**E**) Negative-staining transmission electron microscopy image of MVs purified from the SN of *L. helveticus* VEL12193. (**F**) Size measurement of unlabeled MVs from *L. helveticus* VEL12193 by dynamic light scattering (DLS). (**G**) Representative images of GFP-LC3 HeLa cells treated for 2 h with MVs purified from *L. helveticus* SN or MRS medium (control). LC3 protein coupled to GFP (GFP-LC3) appears in green. (**H**) Quantification of the number of LC3 dots per cell 2h after treatment of GFP-LC3 Hela cells with MVs. Data are mean +/- SEM of three independent experiments. Mann-Whitney test was used and *p*-value is indicated on the graph. (**I**) *In vitro* autophagy flux assay. Representative immunoblots of GFP-LC3 HeLa cells treated or not (control) with MVs purified from the SN of *L. helveticus* (MV *Lh*) for 2 h, in presence of BafA1. Immunoblot analyses were performed using anti-LC3B and anti-actin antibodies. (**J**) The ratio LC3-II/Actin ratio was calculated and normalized to that obtained in control cells without BafA1 (n=5). Wilcoxon matched pairs signed rank test was used.

We hypothesized that membrane vesicles (MVs), which are lipid nanoparticles of high molecular weight able to interact with host cells and modulate their activities, may be involved in the stimulation of autophagy by *L. helveticus* [44,45]. Since to our knowledge there are no reports demonstrating the release of MVs by the bacterial species *L. helveticus*, we first investigated whether *L. helveticus* strain VEL12193 produced such nanoparticles. Analysis of concentrated and purified *L. helveticus* SN revealed the presence of spherical structures with an average diameter of about 150 nm, with most of the vesicles having a diameter from 120 nm to 200 nm (**Figure 5E** and **5F**), a phenotype similar to the MVs produced by other Gram-positive bacteria such as *Lacticaseibacillus casei* [46]. Such structures were not observed in the MRS control (**Sup Figure 3A**).

We next examined the ability of MVs purified from SN *Lh* (MV *Lh*) to stimulate autophagy in GFP-LC3 Hela cells (**Figure 5G-J**). Interestingly, treatment of cells with MV *Lh* induced a significant two-fold increase in the number of GFP-LC3 positive dots, as previously observed with the >100 kDa SN fraction (**Figure 5B, 5G** and **5H**). Although not statistically significant, a trend towards increased LC3-II accumulation was observed in MV *Lh* treated cells compared to control (**Figure 5I** and **5J**).

In order to explore whether the increase in GFP-LC3 positive vacuoles observed in MV *Lh*-treated cells resulted from the endocytosis of these particles into LC3-positive vacuoles - a mechanism termed LC3-associated endocytosis that involves part of the actors of the autophagy machinery [47], we labelled MV *Lh* with a lipophilic probe (DiI) to study their interactions with host cells (**Sup Figure 3B**). We observed that MV *Lh* were rarely (5.6% of total MV *Lh*) found in close proximity of LC3 positive structures (**Sup Figure 3B** and **3C**), suggesting that while MV *Lh* induce autophagy, they are not necessarily taken up extensively within autophagy vacuoles.

Altogether, these results indicate that MVs produced by *L. helveticus* VEL12193 are autophagy inducers.

### Immunomodulatory properties of L. helveticus strain VEL12193 in vitro and in vivo in mice

Since autophagy is intimately linked to the regulation of inflammatory processes [48,49], we evaluated the anti-inflammatory properties of *L. helveticus* strain VEL12193. First, we confirmed *in vitro* that MV *Lh* stimulated autophagy in RAW264.7 macrophages, as shown by a significant increase in the number of LC3 positive vacuoles in MV *Lh _-_*treated macrophages compared to control (**Figure 6A** and **6B**). Next, we analyzed the ability of MV *Lh* to dampen the pro-inflammatory response induced by treatment of RAW264.7 macrophages with lipopolysaccharide (LPS). MV *Lh* treatment of LPS-stimulated macrophages was associated with trends toward decreases of the expression level of the four tested genes encoding the pro-inflammatory cytokines IL-6, TNF-alpha, IL-1beta and IFN-gamma (**Figure 6C-6F**). Analysis of the cytokines released in the supernatant of LPS-treated macrophages showed that MV *Lh* treatment significantly reduced both the secretion of IL-6 and IFN-gamma (**Figure 6G-6I**). To note, IL-1beta was undetectable in all tested conditions. These results suggest that MV *Lh* has anti-inflammatory property.

**Figure 6.**
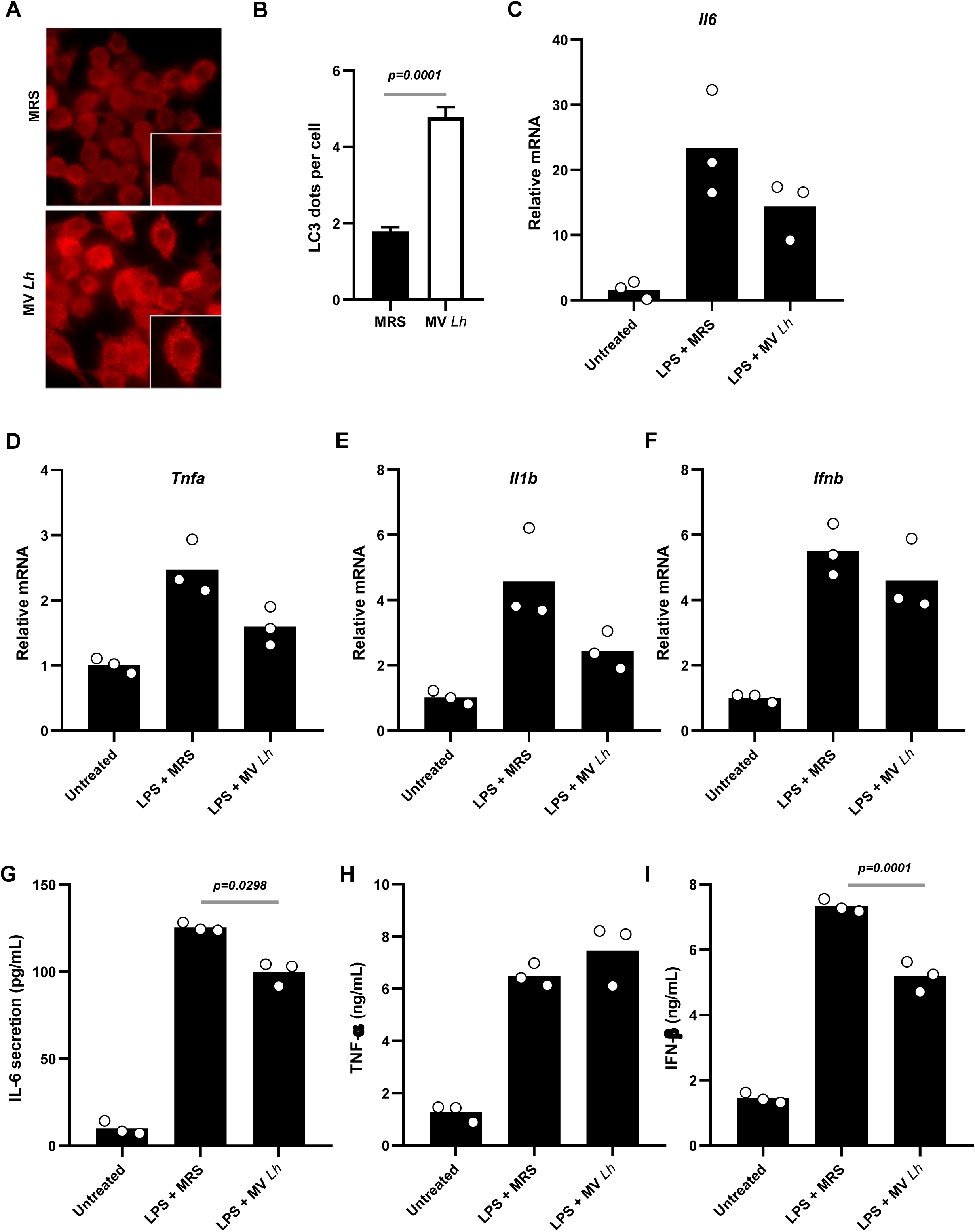
Immunomodulatory properties of L. helveticus-derived membrane vesicles in lipopolysaccharide-stimulated macrophages. **(A**) Representative images of wild-type RAW264.7 macrophages treated for 2h with MVs purified from the SN of *L. helveticus* (MV *Lh*) or bacterial culture medium (MRS medium, control). Cells were immunostained with anti-LC3 antibodies (red). (**B**) Quantification of the number of LC3 dots per cell 2 h after treatment of RAW264.7 macrophages with MVs. Data are mean +/- SEM of three independent experiments. Mann-Whitney test was used and the *p*-value is indicated on the graph. (**C-I**) Wild-type RAW264.7 macrophages were pretreated for 2 h with MV fractions and then stimulated or not with LPS (100 ng/mL) for 4h. (**C-F**) The expression level of genes encoding cytokines (**C**) *Il6*, (**D**) *Tnfa*, (**E**) *Il1b* and (**F**) *Ifnb* was measured by RT-qPCR. The levels of mRNA were normalized to *Hprt* mRNA level for calculation of the relative levels of transcripts. Results are expressed relatively to the expression level of untreated cells. Data are mean ± SEM of three independent experiments. Mann-Whitney test was used, and *p*-values are indicated on the graphs. (**G-I**) The level of (**G**) IL-6, (**H**) TNF-α and (**I**) IFN-β secreted by macrophages (in pg/mL or ng/mL) was determined by ELISA (Mean ± SEM of three independent experiments). Mann-Whitney test was used and *p*-values are indicated on the graphs.

We then investigated whether *L. helveticus* VEL12193 could protect against dextran sulfate sodium (DSS)-induced colitis in mice, a model that reproduce some physiopathological features of human IBDs [50]. For this experiment, we chose to work with living bacteria rather than MV *Lh* to maximize the bioavailability of the active principle in the colon, where the DSS triggered inflammation. Mice were daily supplemented for 15 days with *L. helveticus* and this supplementation was maintained all along the DSS protocol. *L. helveticus* supplementation did not prevent DSS-associated weight loss, a hallmark of colitis severity (**Figure 7A**). The disease activity index (DAI) that includes scores for weight loss, stool consistency and bleeding was calculated. As expected, DSS treatment was associated with a significant increase of the DAI compared to mice of the control group that received PBS glycerol (**Figure 7B**). A trend towards a decrease was observed in the *L. helveticus*-supplemented and DSS-treated group compared to the DSS-treated between day 8 and 10 (**Figure 7B**). After euthanasia, macroscopic examination of the colon showed that *L. helveticus* supplementation did not prevent the reduction of the colon length associated with DSS-induced colitis (**Figure 7C**). Similarly to human IBDs, DSS-induced colitis increased intestinal permeability, a phenotype that can be assessed in mice by measuring the passage of FITC-Dextran in serum after its administration *per os* [51]. We observed that DSS-induced colitis significantly increased intestinal permeability (mean = 14.8±9.9 µg/mL) compared to the control group and that supplementation of DSS-treated mice with *L. helveticus* tended to reduce this phenotype by a two-fold factor (mean = 6.1±4.3 µg/mL) (**Figure 7D**).

**Figure 7.**
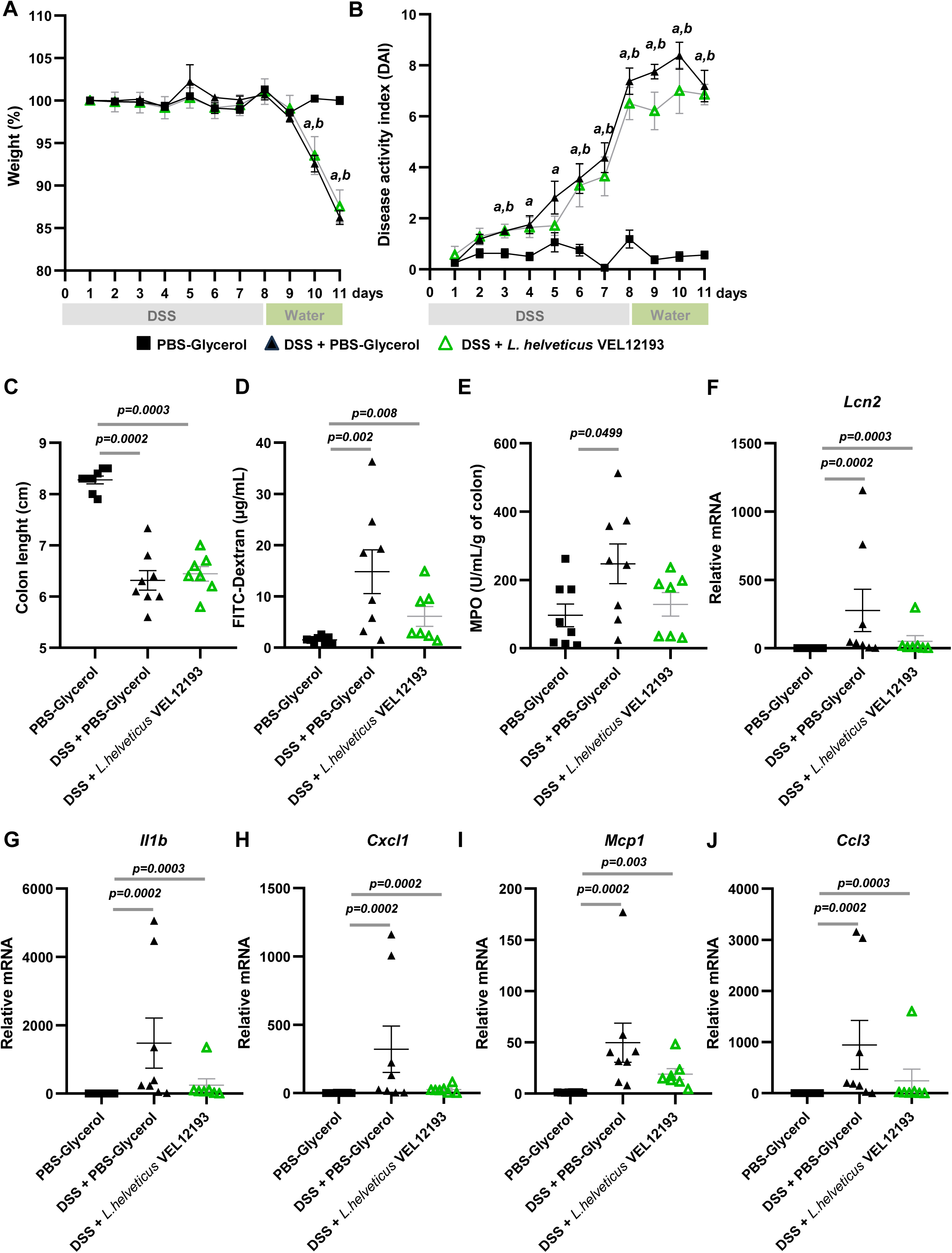
Protective effect of L. helveticus VEL12193 during DSS-induced colitis in mice. Mice were orally gavaged with PBS-Glycerol (control) or *L. helveticus VEL12193* for 15 days before and during the 8 days of DSS-treatment (2% in drinking water) to induce colitis. (D0-D8). Then DSS was removed from drinking water for 3 days (D8-D11, recovery period). Weight (**A**) and disease activity index (DAI, a score that includes weight, bleeding and stool consistency) (**B**) were monitored daily (n=8/group). Results are expressed as mean ± SEM. Data were analyzed by one-way ANOVA followed by Tukey’s multiple comparisons test (p<0.05), a: PBS-Glycerol group versus DSS + PBS-Glycerol group, b: PBS-Glycerol group versus DSS + *L. helveticus* VEL12193, c: DSS + PBS-Glycerol group versus DSS + *L. helveticus* VEL12193. (**C**) Colon length (in cm) expressed as mean ± SEM. (**D**) Intestinal permeability to FITC-dextran (in µg/mL) measured at the endpoint and expressed as mean ± SEM. (**E**) Myeloperoxidase (MPO) activity (in U/mL/g of tissue) measured in colon sections and expressed as mean ± SEM. (**F**-**J**) Expression of genes encoding pro-inflammatory mediators (**F**) *Lcn2*, (**G**) *Il1b*, (**H**) *Cxcl1*, (**I**) *Mcp1* and (**J**) *Ccl3* in mouse colon. The levels of mRNA were normalized to *Hprt* mRNA level for calculation of the relative levels of transcripts. (**C-J**) Mann-Whitney test was used and *p*-values are indicated on the graphs.

Myeloperoxidase (MPO) is an enzyme abundantly produced by neutrophils. Measuring its activity in colon section give a reliable indication of the inflammation severity [51]. As expected, colonic MPO level was significantly higher in DSS-treated mice in comparison to control mice (**Figure 7E**). Mice supplementation with *L. helveticus* prevented this phenotype, the colonic MPO level being not significantly different between the *L. helveticus*-supplemented, DSS-treated group and the control group (**Figure 7E**).

Moreover, we analyzed the colonic expression levels of five genes encoding pro-inflammatory mediators (*Lcn2*, *Il1b*, *Cxcl1, Mcp1 and Ccl3*). DSS-treatment was associated with a significant increase in the mRNA levels of all these inflammatory markers compared to control mice (**Figure 7F-7J**). *L. helveticus* supplementation of DSS-treated mice showed a marked trend towards reduced expression levels of all pro-inflammatory markers (**Figure 7F-J**).

Finally, we evaluated the effect of the DSS-induced colitis on the mRNA expression level of key ATGs genes. Indeed, several studies suggest that some ATGs such as *Atg5* or *Atg7* would play a protective role against DSS-induced colitis [52,53]. We observed that DSS-induced colitis triggered a significant decrease in the colonic expression level of *Atg5*, *Atg7*, *Atg12*, *Becn1*, *Ulk1* and *Vps34*, and a significant increase in the colonic expression level of *Atg16l1* compared to control mice (**Sup Figure 4**). *L. helveticus* supplementation enabled to significantly attenuate the decrease in the colonic expression level of *Atg12* associated to DSS treatment (**Supp Figure 4D**). A similar trend, but not significant, was observed for *Atg5* expression (**Supp Figure 4A**).

Taken together, these results – some significant, others trendy – suggest that *L. helveticus* may attenuate inflammation and that this could involve MV *Lh*. Role of autophagy modulation by *L. helveticus* in such phenotype remain unclear.

## Discussion

Autophagy machinery crucial to maintain homeostasis throughout life in eukaryotic organisms by modulating key cellular processes such as membrane dynamics, removal of damaged organelles, maintenance of energy balance or defense against pathogens [2,54]. Stimulating autophagy has been shown to be beneficial in various physiopathological contexts and to support healthy aging [4]. Various regulatory pathways of autophagy are sensitive to microorganisms and their metabolites, as evidenced by the gut microbiota’s reported modulatory effects on host autophagy [25]. In this study, we explored the potential of food grade bacteria and bacteria belonging to species used as probiotics to stimulate autophagy by combining *in vitro* and *in vivo* approaches.

A growing number of studies described the ability of specific strains from the lactobacilli family or the *Bifidobacterium* genus to modulate host autophagy *in vitro* [55–57]. In this study, we showed that not all strains of lactobacilli and bifidobacteria were able to stimulate autophagy and that it is important to verify this phenotype by different complementary approaches. Indeed, among the 11 strains we tested only one showed a robust pro-autophagy property that was validated by the different approaches used. To date, only a few studies have analyzed *in vivo* the ability of beneficial microorganisms to modulate autophagy in mammals [55,58–60]. Indeed, even if many autophagy-related molecular markers exist, it remains highly challenging to properly assess autophagy responses *in vivo*, notably due to the highly dynamic nature of this process [61]. In addition, autophagy analysis *in vivo* is in general limited to the analysis of the basal expression level of ATGs proteins or genes, which are parameters that do not fully inform on the activity status of this process. In the present study, we validated the *in vivo* pro-autophagic property of *L. helveticus* VEL12193 by combining again different approaches including a study of autophagy flux on *ex vivo* tissue. We chose to work *ex vivo* rather than *in vivo* - which is possible by injecting autophagy inhibitors to mice - for several reasons. From the point of view of robustness of the results, each individual constitutes its own control, thus limiting the variability linked (i) to the use of two different animals, one serving for the analysis of autophagic flux in “open” flux (without the use of autophagic flow inhibitors) and the other for the analysis of autophagic flux in “closed” flux (after injection of autophagic flow inhibitors), and (ii) to the bioavailability of autophagic flow inhibitors for the organs. *Ex vivo* analysis of autophagy flux enables also to limit the number of mice used in the experiment (each animal being its own control). Some tools are available to study autophagy flux in tiny model organisms such as in *Caenorhabditis elegans* or in Zebrafish [62,63] but there is currently no equivalent in rodents.

The prototypical ATG protein LC3 is widely used as a gold standard autophagy marker, since its conjugation to membrane forming autophagosome is a key event during canonical autophagy. However, this protein can also be conjugated to other cellular membranes, a recently described process termed ATG8ylation [64]. ATG8ylation involved some ATG proteins of canonical autophagy such as ATG16L1 but not all. In particular, WIPI2 and ATG13 are not involved. Colocalization of WIPI2 protein with *LC*3 puncta in cells treated with *L. helveticus* and MV *Lh* suggest that this bacterium stimulates canonical form of autophagy. However, it is important to remain cautious in drawing conclusions and not to exclude the possibility of non-canonical forms of autophagy, which may involve only parts of the autophagy machinery without the formation of canonical double-membraned autophagosomes.

Selective autophagy corresponds to the lysosomal degradation of specific intracellular components (e.g., proteins aggregate (aggrephagy), mitochondria (mitophagy), intracellular bacteria (xenophagy), peroxisome (pexophagy)). This involves activity of selective autophagy receptors (SARs) which interact with ATG8 family proteins [65]. Impairment of certain forms of selective autophagy has been associated with the development of pathophysiological conditions such as xenophagy in IBD such as Crohn’s disease, mitophagy and/or aggrephagy in age-related diseases such as AMD and Alzheimer’s disease [66]. Thus, it might be of interest to go further in the analysis of pro-autophagy property of *L. helveticus* and its related MVs by analyzing selective forms of autophagy, which can be done by using specific markers of these processes [65]. By the way, it has been reported that the yeast *Saccharomyces boulardii* strain CNCM-I-1079 and the bacteria *Lactococcus lactis* strain R1058 have the ability to stimulate mitophagy *in vitro* and *in vivo* in drosophila model, thus leading to neuroprotective effects [55].

In our study, we observed that *L. helveticus* VEL12193 can mitigate some of the traits of DSS-induced colitis (e.g., intestinal permeability, neutrophils infiltration or the expression of pro-inflammatory cytokines). It would also be interesting to evaluate the pro-autophagic activity of the *L. helveticus* strain in other physiopathological conditions that challenge the autophagy pathway locally in the gut, such as infection by enteric pathogens [67] or at distance in models of degenerative diseases [68]. Symptoms alleviation by bacterial strains of the lactobacilli family with pro-autophagic property have been reported in mice infected with *Salmonella enterica* Typhimurium) or after TNBS-induced colitis, but no causality link between the stimulation of autophagy and symptom alleviation has been demonstrated [57,58]. This is also one limitation in our study. Further experiments using adult mice with conditional knockout of genes encoding key ATG proteins in intestinal epithelial or immune cells could help to determine whether this protective effect during colitis is dependent on autophagy. But it should be kept in mind that such experiments carry a risk of misinterpretation. Indeed, the autophagy machinery is more complex than previously thought, and knockout mice for different key ATG proteins exhibit varying phenotypes [69].

Even if a growing number of studies point out the ability of some beneficial microbes to stimulate autophagy, to our knowledge, no study has yet definitively identified the specific microbial compound(s) responsible for this pro-autophagic property. Recurrently in studies, the supernatant of these bacteria, including strains of *Levilactobacillus brevis*, *Lactiplantibacillus plantarum*, *Lacticaseibacillus rhamnosus* and *Lacticaseibacillus paracasei,* has been identify as the active fraction [58,70–73]. In our study, we identified the MVs released in the supernatant of *L. helveticus* as an active compound contributing to autophagy stimulation. It might be of interest to analyze whether MVs are also responsible for the supernatant-induced autophagy stimulation described for other strains. Differently to Gram-negative bacteria in which MVs formation occur directly through outer membrane blebbing giving outer-membrane vesicles (OMVs), the formation of MVs in Gram-positive bacteria, such as *L. helveticus*, requires local or global (bubbling cell death) peptidoglycan degradation, allowing cytoplasmic membrane to protrude through holes in the peptidoglycan layer [74]. In comparison to the production of MVs by Gram-negative bacteria, far less is known regarding the regulation of MVs production by Gram-positive bacteria. Several stress such as antibiotics treatment (β-lactam family), DNA damaging agents or peptidoglycan degrading enzymes produced by other microorganisms that weaken the bacterial cell wall have been identified to enhance MVs production in Gram positive bacteria [46,75]. Physiological stimuli promoting MVs production in Gram positive bacteria, notably in the context of the gastrointestinal tract environment, remain to be determined. It would be particularly interesting to investigate the *in vivo* production of these MVs in the gastrointestinal tract, where they can interact with intestinal epithelial cells and potentially diffuse to peripheral organs, as suggested for some Gram negative bacteria of the gut microbiota [76].

Analyses of the composition of MVs released by Gram positive bacteria have shown that it is highly complex and includes lipids (e.g., phospholipids or lipoteichoic acid), hundreds of proteins, nucleic acids and other metabolites [75]. This complexity makes it challenging to pinpoint the specific compound(s) responsible for an effect, in our case stimulation of autophagy in host cells. DNA, RNA and peptidoglycan of MVs derived from *Staphylococcus aureus* have been reported to activate in lung epithelial cells Toll-like receptor (TLR) 2, 7, 8 and 9 as well as the intracellular Nucleotide-binding Oligomerization Domain 2 (NOD2) receptor [77]. Activation of these receptors, alone or in combination, have been shown to modulate autophagy in various cell types [22,77]. One could assume that such bacterial molecular pattern and host receptor could also contribute to autophagy stimulation by *L. helveticus* strain VEL12193 [78].

It has been proposed that OMVs released by Gram negative bacteria can be internalized in non-phagocytic host cells through several routes including clathrin- or caveolin-mediated endocytosis, micropinocytosis or lipid rafts [79]. Fusion events of MVs from Gram-positive pathogens such as *S. aureus* or *Listeria monocytogenes* with host cells have also been suggested, enabling the transfer of virulence factors (α-Hemolysin and Listeriolysin O, respectively) to the host plasma membrane [80,81]. Far less is known about the trafficking of MVs from beneficial bacteria within host cell. Endocytosis of MVs released by *Lacticaseibacillus rhamnosus* having immunomodulatory effects has been described *in vitro* in intestinal epithelial cells [82]. We observed that *L. helveticus* MVs were tightly associated with host cell and that some of them colocalized within LC3 positive structures, offering the intriguing possibility that internalized bacterial MVs can be handled into endomembrane connected to the autophagy pathway. This has already been suggested for OMVs released by the Gram-negative bacteria *Helicobacter pylori* and *Pseudomonas aeruginosa* that colocalized with endosomes, in close association with LC3-positive vacuoles [83]. Similarly, *S. aureus*-derived MVs was shown to colocalize with LC3-positive structures [77]. Further investigations are needed to better understand the molecular interactions between *L. helveticus* MVs and the autophagy machinery.

The *L. helveticus* strain VEL12193 identified in this study has been the best candidate among lactobacilli and bifidobacterial tested to stimulate autophagy in host cells has been isolated from a fermented food, a French cheese (Comté). This reinforces the idea that fermented foods represent a tremendous source for discovering microorganisms with significant potential benefits for human health. Indeed, there are an estimated 7,500 species of fungi and bacteria associated with fermented foods, each comprising many different strains. [84]. The abundance and bioavailability of the bioactive metabolites produced by these microorganisms would depend on their local environment (e.g., plant, food matrix or digestive tract). Food matrices can modify probiotic functionalities in term of survival in the gastrointestinal tract or ability to colonize the gut mucosa [85]. For instance, some lactic acid bacteria isolated from traditional fermented foods (cheese, kimchi, or fermented soybean) can produce Gamma-aminobutyric acid (GABA), an amino acid with multiple physiological functions, including autophagy stimulation in macrophages and its associated antimicrobial response [86], and its production is strain- and food-matrices-dependent [87]. Indeed, the production of GABA is controlled by the glutamate decarboxylase (GAD), and highly dependent on the presence of the precursor L-glutamic acid and the cofactor pyridoxal 5′-phosphate in the bacterial microenvironment. Moreover, production of bioactive molecules by microorganisms can also be modulated by diet. In a human trial, consumption during 5 days of a plant-based diet was sufficient to induce a two-fold increase in the production of the SCFA acetate and butyrate by the gut microbiota in comparison to the level observed in individuals consuming an animal-based diet [88]. Given the important role of SCFA in the regulation of autophagy [25], we can hypothesize that such diet-induced changes of SCFA abundance can potentially impact autophagy activities in host cells.

To meet the major public health challenge facing our societies as the population ages and the prevalence of chronic diseases rises, sustainable and safe preventive solutions need to be developed. Consumption of fermented foods is associated with various health benefits - including anti-aging related benefits - that are mediated by the microorganisms, and their derived products [84,89] and contribute to sustainable development by promoting efficient resource use and minimizing environmental impact, aligning seamlessly with the ‘One Health’ approach. Explore the microbial biodiversity offered by fermented foods and better characterized how they can modulate key host cellular processes is essential to unlock their full potential and enable the development of new microbial-based foods that support whole-body homeostasis and promote healthy aging.

To conclude, in this study, we underscore the potential of the food-grade bacteria *L. helveticus* and its derived products (MVs) to promote autophagy *in vitro* and *in vivo*. Further investigations are now required to explore in-depth the interaction of bacterial MVs with the host cells and their potential intracellular trafficking. In addition to the fundamental insights provided by these results, they represent proof-of-concept for the research and development of innovative microbial-based functional foods (probiotics and postbiotics) aimed at promoting the cytoprotective mechanism of autophagy.

## Material and methods

### Bacterial strains and culture conditions

Bacterial strains used in this study are described in **Table 1**. Bacteria were grown routinely under anaerobic conditions at 37□°C without shaking in Man-Rogosa-Sharpe (MRS) medium (Condalab), buffered at pH 5.8.

### Cell lines and treatment with bacteria

All cell lines were maintained in an atmosphere containing 5% CO_2_ at 37°C and were routinely tested for mycoplasma contamination using the PCR Mycoplasma Test Kit II (PromoKine). The human colonic epithelial cell line HCT116 was obtained from ATCC. The human epithelial HeLa cells stably expressing GFP-LC3 were generously donated by Mathias Faure (Inserm U1111, CIRI, Lyon, France). All epithelial cells were maintained in DMEM-GlutaMax (Gibco) supplemented with 10% (vol/vol) fetal bovine serum (FBS, PanBiotech) without antibiotics. The murine macrophage cell line RAW 264.7 was obtained from ATCC and maintained in RPMI medium 1640 (Gibco) supplemented with 10% (vol/vol) FBS (PanBiotech) without antibiotics.

For treatment with bacteria, cells were seeded into 24-well culture plates at a density of 2.10^5^ cells per well and incubated for 48 h. Cells were washed twice with phosphate-buffered saline (PBS; pH 7.2), and the medium was replaced by 1 ml of DMEM-GlutaMax supplemented with 10% heat-inactivated FCS. Epithelial cells were treated at a ratio of 10 bacteria per cell for 2 h or 4 h.

### Antibodies and staining reagents

Inhibitors of autophagy flux used in the study were bafilomycin A1 (Baf A1, Merck **#**B1793), leupeptin (Merck, **#**L2884) and NH_4_Cl (Merck**, #**A9434). Lipopolysaccharides (LPS) from *Escherichia coli* O127:B8 (Merck, #L3129) was used to stimulate inflammatory response.

For western blot analysis, the primary antibodies used were anti-LC3B (Merck, **#** L7543), anti-SQSTM1 (p62; Abnova **#**H00008878-M-01), anti-Lamp2a (Abcam, **#**ab18528) and anti-Actin (Merck, **#**A2066). The secondary antibodies used were polyclonal Goat Anti-Rabbit Immunoglobulins/HRP (Agilent, **#**P044801-2) and polyclonal Goat Anti-Mouse Immunoglobulins/HRP (Agilent, **#**P044701-2).

For immunostaining, the primary antibodies used were anti-LC3 (MBL, **#**PM036) and anti-WIPI2 (Millipore, **#**MABC91). Nuclei were stained using DAPI (Merck, **#**D9542).

### Immunofluorescence

For *in vitro* experiments, cells were seeded on glass coverslips in 24-well plates. Briefly, at the end of the experiment, cells were washed with PBS, fixed with PBS-4 % paraformaldehyde (PFA) for 10 min, and then permeabilized and saturated in PBS-3% BSA-0.05% Triton X-100 for 20 min. Cells were then incubated at room temperature (RT) for 2 h with primary antibodies diluted in PBS-3% BSA, washed and incubated for 1 h at RT with Alexa-fluor conjugated secondary antibodies and DAPI diluted in PBS-3% BSA. Images were acquired using Axiovision Zeiss fluorescent microscope. The number of LC3 or WIPI2 dots per cell were counted in at least 100 cells for each experiment using Icy software [96]. Each microscopy image is representative of at least three independent experiments.

For mouse colon samples, tissues were fixed in 10% buffered formalin, embedded in paraffin, and sectioned. Sections were dewaxed and hydrated by incubating slides successively in xylene, xylene/ethanol, ethanol (100%-50%) and water baths. Antigen retrieval was performed enzymatically by incubating slides in a pre-warmed (37°C) 0.05% trypsin, 1% NaCl water solution for 15 min. After washing, sections were incubated at RT in TBS-10% FBS-1% BSA for 2 h. Then, sections were incubated overnight at 4°C with primary antibody, washed twice with TBS-0.025% Tween-20, and incubated with Alexa-fluor conjugated secondary antibody and DAPI in TBS-1% BSA for 1 h at RT. Images were acquired using Axiovision Zeiss fluorescent microscope. The mean intensity of LC3 staining in the gut mucosa was determined using ImageJ software on colon samples from four mice for each group, with measurement over at least 5 fields for each mouse.

### Autophagy flux assay

#### In vitro experiments

Bafilomycin A1 (BafA1) was used to inhibit autophay flux *in vitro*. The BafA1 was added at 100 nM in the cell culture medium used during bacteria/cell interaction experiments.

#### Ex vivo experiments

Autophagy flux was assessed *ex vivo* on dissected retina and colon. Retinas from a same mouse were placed into two independent wells of a 96-well cell culture plate. One retina was incubated for 4h in DMEM 4.5 g/L glucose Glutamax® (Pan-Biotech), containing 10% fetal calf serum (Biosera) and 10 µg/mL antibiotics (penicillin and streptomycin, Pan-Biotech) while the other retina was incubated for 4 h in the same medium supplemented with 100 µM leupeptin (Merck, L2884) and 20 mM NH_4_Cl (Merck).

For each mouse, the colon was opened longitudinally, and two adjacent areas of the same size were cut out and deposited (intestinal mucosa exposed upwards) into two independent wells on a 96-well cell culture plate. As described for retinas, one of the two pieces of colon was incubated for 4h in complete cell culture medium without autophagy inhibitors while the other piece was incubated for 4 h in the same medium supplemented with autophagy inhibitors (100 µM leupeptin and 20 mM NH_4_Cl).

At the end of the 4 h incubation period, colon and retina samples were snap-frozen in liquid nitrogen and store dried at −80°C until Western blot analyses.

### Western blot

Whole-cell protein extracts were prepared from cultured cells by adding directly 200 µL of 1.25X Laemmli sample buffer on the cell monolayers, and by incubating the lysates for 5 min at 95°C. Whole-cell protein extracts were prepared from retina and colon samples by using RIPA lysis buffer (Fisher) supplemented with protease and phosphatase inhibitors (PhosSTOP^TM^ and cOmplete^TM^ ULTRA tablets, Merck). Proteins were separated on 4–15% precast polyacrylamide gels (Bio-rad), transferred on to nitrocellulose membrane (BioRad), blocked for 1 h in Tris-buffered saline (TBS) solution containing 5% non-fat dry milk, and probed overnight with primary antibodies and for 2h with secondary HRP-coupled antibodies. Actin level was used to normalize protein quantity. After membrane revelation using ECL detection kit (Perkin Elmer), quantification was performed with ImageJ software.

### RT-qPCR and qPCR

RNAs were extracted and purified from samples using NucleoSpin RNA/Protein kit (Macherey-Nagel). Reverse transcription was performed with PrimeScript RT reagent kit containing gDNA Eraser (Takara Bio Europe) and using 500 ng of total RNA. Gene expression was determined by real-time polymerase chain reaction using SYBR Green (Bio-Rad) and a StepOnePlusTM Real-Time PCR System (Fisher Scientific). *Hprt* was used as the internal control for normalization. Fold induction was calculated with the delta-delta Ct (ddCt) method. Primer sequences are given in **Table 2**.

**Table 2.**
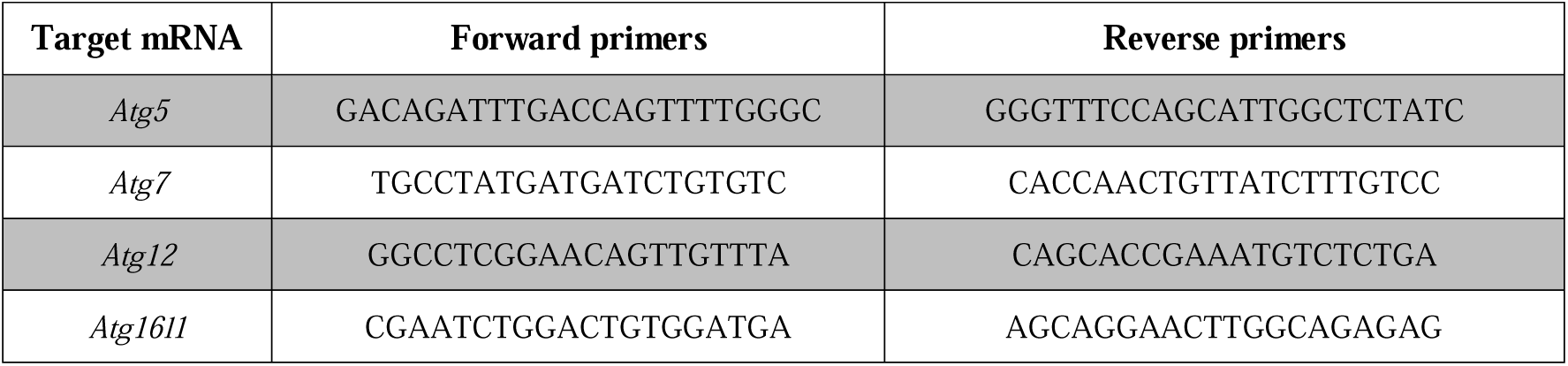
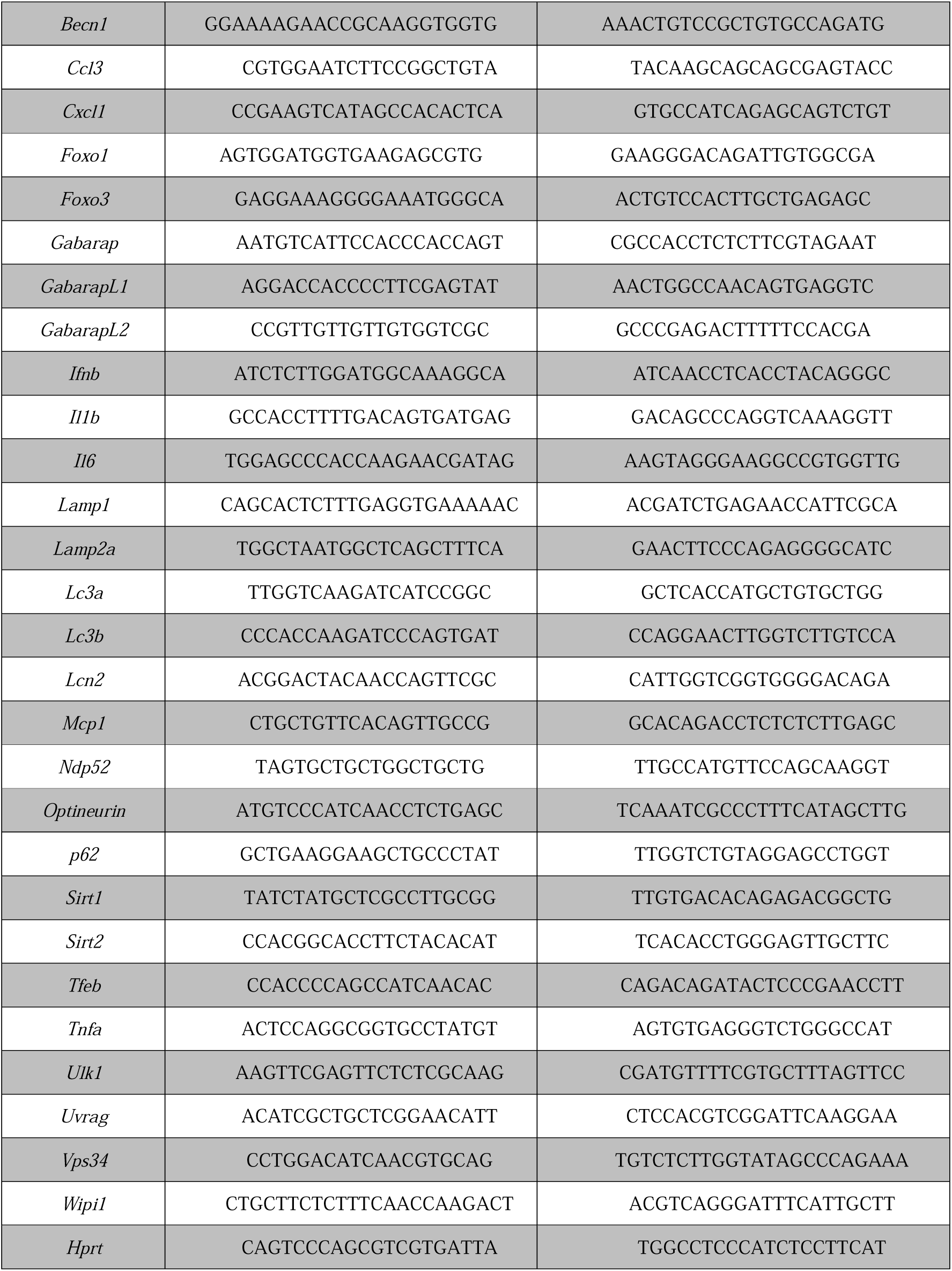
Murine primers used in this study.

For the quantification of *L. helveticus*, 200 ng of fresh feces were collected. Total DNA was extracted and purified using ZymoBIOMICS DNA Miniprep kit (Zymo Research). The quantity of *L. helveticus* DNA was determined by qPCR analysis using iTaq universal SYBR Green supermix (Bio-Rad) and the following primers: forward 5’-AGGTTCAAAGCATCCAATCAATATT-3’ and reverse 5’-TCGGGACCTTGCACTACTTTATAAC-3’. Fold changes in *L. helveticus* DNA were calculated using the ΔΔCt method of relative quantification, with the total bacteria as a reference using the following primers: forward” 5’-CAGCAGCCGCGGTAATAC-3’ and reverse 5’-CCGTCAATTCCTTTGAGTTT-3’.

### Mice

All animal experiments were performed following European Union guidelines and the ARVO Statement for the Use of Animals in Ophthalmic and Vision Research. French legal and institutional ethics committee review board approvals were obtained (2018072513005644). Twelve-month-old male C57BL/6JRj SPF mice were purchased from Janvier Labs, France. They were maintained at INRAE, Dijon, France until euthanasia (C21 231 010 EA) with *ad libitum* access to food and water and exposed to 12□h:12□h light:dark cycles. After one week of acclimation, mice were randomly divided into two groups: one group received standard diet (control group; n□=□10) and the other group received the same diet as controls but supplemented with *L. helveticus* (*L. helveticus* group; n□=□10). Mice were maintained on these diets for 6 months. Prior to euthanasia, mice were fasted for 15□h to stimulate basal autophagy flux. They were euthanized by cervical dislocation. Colon and retina were collected.

### Diet

*L. helveticus* strain VEL12193 (**Table 1**) was grown overnight under anaerobic conditions at 37□°C without shaking in MRS medium, pH 5.8. The bacterial culture was centrifuged at 5000□g for 10□min at RT. The bacterial pellet was washed twice in PBS and resuspended in sterile water at a concentration of 2.10^9^ CFUs/mL. This bacterial suspension was then mixed with complete maintenance diet powder for adult mice (SAFE® A04) to obtain a final bacterial concentration in the diet of 1.10^9^ CFUs/g. Food portions (approximatively 20□g) were molded into Petri dishes, dried for 24□h at 4□°C and then stored anaerobically at 4□°C. Fresh diet was prepared weekly. The food portions were renewed in the cages every 2 days from the stock stored in anaerobic conditions at 4□°C. The viability of *L. helveticus* in food portions stored under these conditions was checked by the agar plate (MRS) method [33].

### Fractionation of bacterial cell free supernatant (SN) and purification of membrane vesicles (MV)

A protocol adapted from a previous study was used [97]. A culture of *L. helveticus* (250 mL) was centrifuged at 4000 x□g and 4 °C for 20 min. The SN was filtered through a 0.45 µm pore size filter (Nalgen Rapid-Flow) to remove remaining bacterial cells and the pH was adjusted to 7.0 with NaOH. Next, the SN was concentrated by ultrafiltration at 4000□g and 4°C using 100 K Amicon® Ultra-15 centrifugal filters (Merck Millipore). The concentrated SN was filtered through a 0.45-µm-pore-size filter. This fraction was the SN *L. helveticus* >100kDa (SN *Lh* >100 kDa). The sample was then ultracentrifugated at 125,000 g and 4 °C for 2 h. The pellet of MVs was resuspended in 400 µL of sterile PBS (1X). This fraction was the *L. helveticus* MVs (MV *Lh*). The protocol to obtain the fraction SN >3kDa of *L. helveticus* (SN *Lh* >3 kDa) was similar as the one described above at the exeption that the SN was concentrated by ultrafiltration at 4000□g, 4°C with 3 K Amicon® Ultra-15 centrifugal filters (Merck Millipore). All fractions were stored as aliquots at − 80 °C for later use. To constitute controls, the same procedures were applied to the MRS culture medium for each fraction. For experiment, fractions were used at a final concentration of 10% (vol/vol) in the cell culture medium.

### Transmission electron microscopy

Five microliters of sample (MV *Lh* or MRS) were applied and incubated for 1 min to glow-discharged, carbon-coated formvar 200-mesh grids. Grids were blotted and then stained with Uless solution ® (Delta microscopies, USA) for 2 min. Visualizations were performed at the DimaCell platform (http://www.dimacell.fr/) using a transmission electron microscope (HITACHI HT7800) operating at 80 kV and equipped with two AMT cameras (NanoSprint43 AMT, Woburn, USA).

### Dynamic Light Scattering (DLS) measurement

Dynamic light scattering (DLS) measurements were carried out using Submicron Particle Sizer NICOMP 380. MVs were loaded into disposable cells, and data were collected at 25 °C. For each sample, the autocorrelation function was an average of three runs of 2 minutes each and then repeated three times.

### MVs staining to monitor their uptake and trafficking in host cells

MVs were purified as described above at the exception that the pellet was resuspended, diluted 10 times and stained with 1% Vybrant DiI cell-labelling (Invitrogen, V22889) at 37°C for 30 min. The solution is then added on the top of a gradient (OptiPrep™, D1556) and Tris-sucrose buffer solution (0.25 M sucrose, 10.02 µM Tris-HCl, pH 7.0) and ultracentrifugated at 100,000 g and 4°C for 18 h. The stained MVs were collected between the layers 20%-40% of iodixanol, resuspended in PBS and ultracentrifugated at 125,000 g and 4°C for 2 h. Finally, the pellet was resuspended in 200 µL PBS. As a control, the same procedure was applied to the purification product made on growth medium (MRS). Purified stained MVs were added at a final concentration of 5% (vol/vol) to HeLa-GFP LC3 cells, followed by a centrifugation at 200 g for 10 min. The cells have been observed by fluorescent microscopy after a 6 h incubation with MVs at 37°C.

### Treatment of cells with lipopolysaccharide (LPS)

WT RAW264.7 macrophages were treated 2 h before LPS stimulation by *L. helveticus* MVs or the control fraction (MRS). LPS stimulation was achieved by adding LPS at a final concentration of 100 ng/mL for 4 h. Cell supernatants were collected for ELISA and cell monolayer harvested for mRNA extraction.

### ELISA

Cytokine secretion by WT RAW 264.7 macrophages stimulated for 4 h with LPS was measured by ELISA using BD OptEIA™ Mouse IL-6 ELISA Kit (BD Biosciences), BD OptEIA™ Mouse TNF ELISA Kit (BD Biosciences) and Mouse IFN-beta DuoSet ELISA (R&D Systems) according to the manufacturers’ instructions.

### Dextran Sulfate Sodium (DSS)-induced acute colitis in mice

The protocol used in this study was based on those published previously [98]. Prior to colitis induction, mice were gavaged with 1 × 10^9^ CFU of *L. helveticus* strain VEL12193 suspended in 200 μL of PBS or with 200 μL of PBS alone, daily for 15 days. Acute colitis was induced by exposing mice for 8 days to drinking water containing 2% (weight/vol) DSS with a molecular weight of 36,000-50,000 kDa (MPBio). Control mice received standard drinking water. On day 8, DSS-treated mice were put back on standard drinking water for 3 days (recovery period). On day 11, the mice were euthanized by cervical dislocation. During the experiment, mice were monitored daily for weight loss, stool consistency and the presence of blood in the stool (Hemoccult, Beckman Coulter). The disease activity index (DAI) was calculated according to the protocol established by Cooper and colleagues [99].

### Intestinal permeability *in vivo*

Intestinal permeability was assessed *in vivo* using fluorescein isothiocyanate-conjugated dextran (FITC-dextran 3000-5000 Da, Merck) as a tracer, as previously described [100]. Briefly, at the end of the experiment, FITC-dextran was dissolved at 0.6 mg per g of body weight in PBS, and this solution was administered to mice by oral gavage. To measure the passage of FITC-dextran into the blood, blood samples were taken 3.5 h after gavage from the retro-orbital venous plexus. These samples were stored in the dark at 4°C until analysis. For the assay, serum was separated by centrifugation and FITC levels in plasma were determined using a fluorescence microplate reader (excitation 485 nm and emission 530 nm; Tecan).

### Myeloperoxidase (MPO) activity

MPO activity was measured according to the method of Bradley et al. [101], modified as described below. One-centimeter length of colon was homogenized (50 mg/mL) in ice-cold 50mM potassium phosphate buffer (pH 6) containing 5% hexadecyl trimethyl ammonium bromide (Merck) and hydrogen peroxide (0.0005%). The colorimetric reaction was followed by measuring the absorbance with a spectrophotometer (460 nm). MPO activity was expressed as units per milligram of wet tissue.

### Statistical analyses

Statistical analyses were performed using Prism 9 software (GraphPad Software Inc.). ANOVA one way, Wilcoxon, Kruskal-Wallis and Mann-Whitney tests were used in this study. All *p*-values of less than 0.05 were considered statistically significant.

## Acknowledgments

This work was funded by Agence Nationale de la Recherche [ANR-11-LABX-0021-01]; INRAE; French “Investissements d’Avenir” program, project ISITE-BFC (contract ANR-15-IDEX-0003); Conseil Régional de Bourgogne, Franche-Comté [PARI grant]; FEDER (European Regional Development Fund) and Institut Carnot Qualiment [grant INPROBIAUS]. We thank Laure Avoscan for TEM experiments (Dimacell Microscopy Facility)

## Data availability statement

The data that support the findings of this study are available from the corresponding authors upon reasonable request.

## Legends of supplementary figures

***Supplementary Figure 1. Colocalization of LC3 and WIPI2 in L. helveticus-treated GFP-LC3 Hela cells.*** Representative images of GFP-LC3 HeLa cells treated for 2 h with *L. helveticus* VEL12193. Cells were immunostained with antiWIPI2 antibodies (red). LC3 protein coupled to GFP (GFP-LC3) appears in green. White squares in upper panels indicate inset areas displayed in the corresponding lower panels.

***Supplementary figure 2: Long term administration of L. helveticus VEL12193 in mice.*** (A) Fecal presence of *L. helveticus* measured by qPCR in mice fed a control diet (black dots) or fed a diet supplemented with *L. helveticus* VEL12193 (white dots) (n=4). The level of *L. helveticus* DNA was normalized to Total bacteria DNA level. (**B**) Weight follow-up during 6 months of mice fed a control diet (black squares, n=9) or fed a diet supplemented with *L. helveticus* VEL12193 (white squares, n=8). Mann-Whitney test was used and p-values are indicated on the graphs.

***Supplementary Figure 3: Microscopic analyses of membrane vesicles released by L. helveticus VEL12193.*** (**A**) Negative-staining transmission electron microscopy image of purified MV fractions obtained from bacterial cell culture medium (MRS medium, control) or SN of *L. helveticus* VEL12193. (**B**) Representative images of GFP-LC3 HeLa cells treated for 6h with Vybrant DiI-labelled purified MV fractions from MRS medium (control) or SN of *L. helveticus* VEL12193. MVs labelled with DiI are in red and LC3 protein coupled to GFP (GFP-LC3) appears in green. White squares indicate inset areas. (**C**) Percentage of *L. helveticus*-derivedMVs (DiI-positive) co-localizing with GFP-LC3 positive structures (dots).

***Supplementary Figure 4: Expression of ATG genes in the colon of mice with DSS-induced colitis.*** (**A-H**) Expression of autophagy-related genes (**A**) *Atg5*, (**B**) *Atg7*, (**C**) *Atg12*, (**D**) *Atg16l1*, (**E**) *Becn1*, (**F**) *Tfeb*, (**G**) *Ulk1*, (**H**) *Vps34* in the colon of mice gavaged with PBS-Glycerol or *L. helveticus* VEL12193 and receiving or not DSS 2% in the drinking water. PBS-Glycerol group: n=8. DSS + PBS-Glycerol group: n=8. DSS + *L. helveticus* VEL12193 group: n=7. Mann-Whitney test was used and *p*-values are indicated on the graphs (NS: non-significant).

