## Supplementary figures and images for "The food grade bacterium *Lactobacillus helveticus VEL12193* promotes autophagy by releasing membrane vesicles"

### Supplementary figures 1 to 4

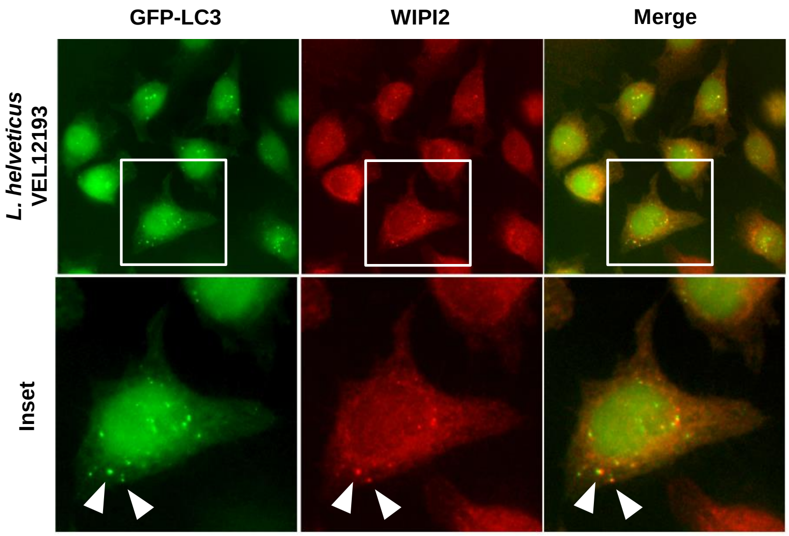


#### Supplementary figure 1

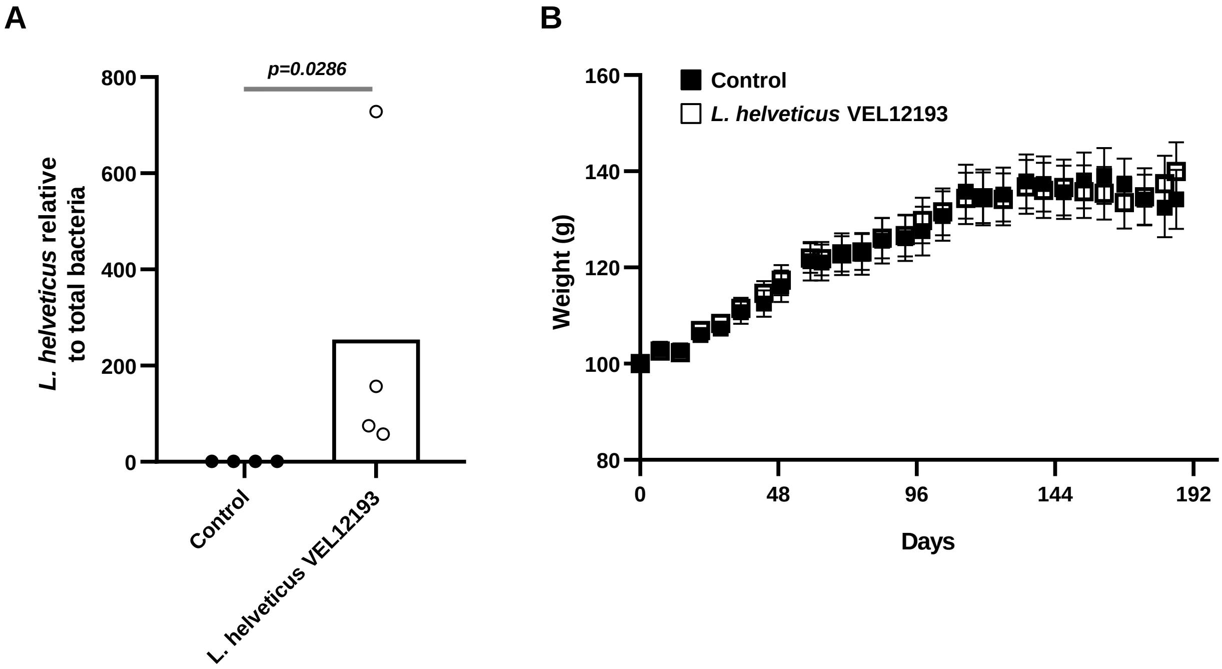


#### Supplementary figure 2

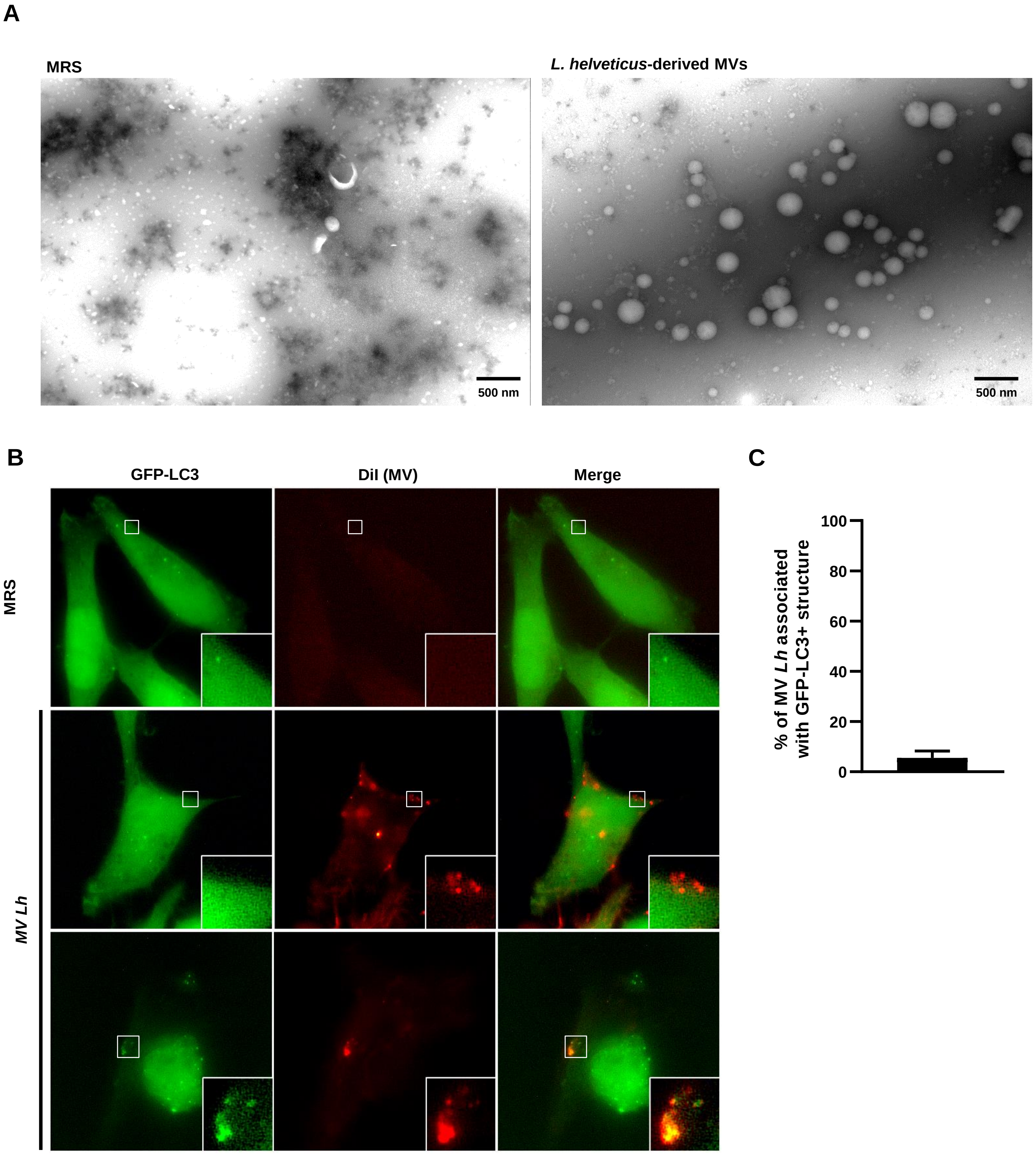


#### Supplementary figure 3

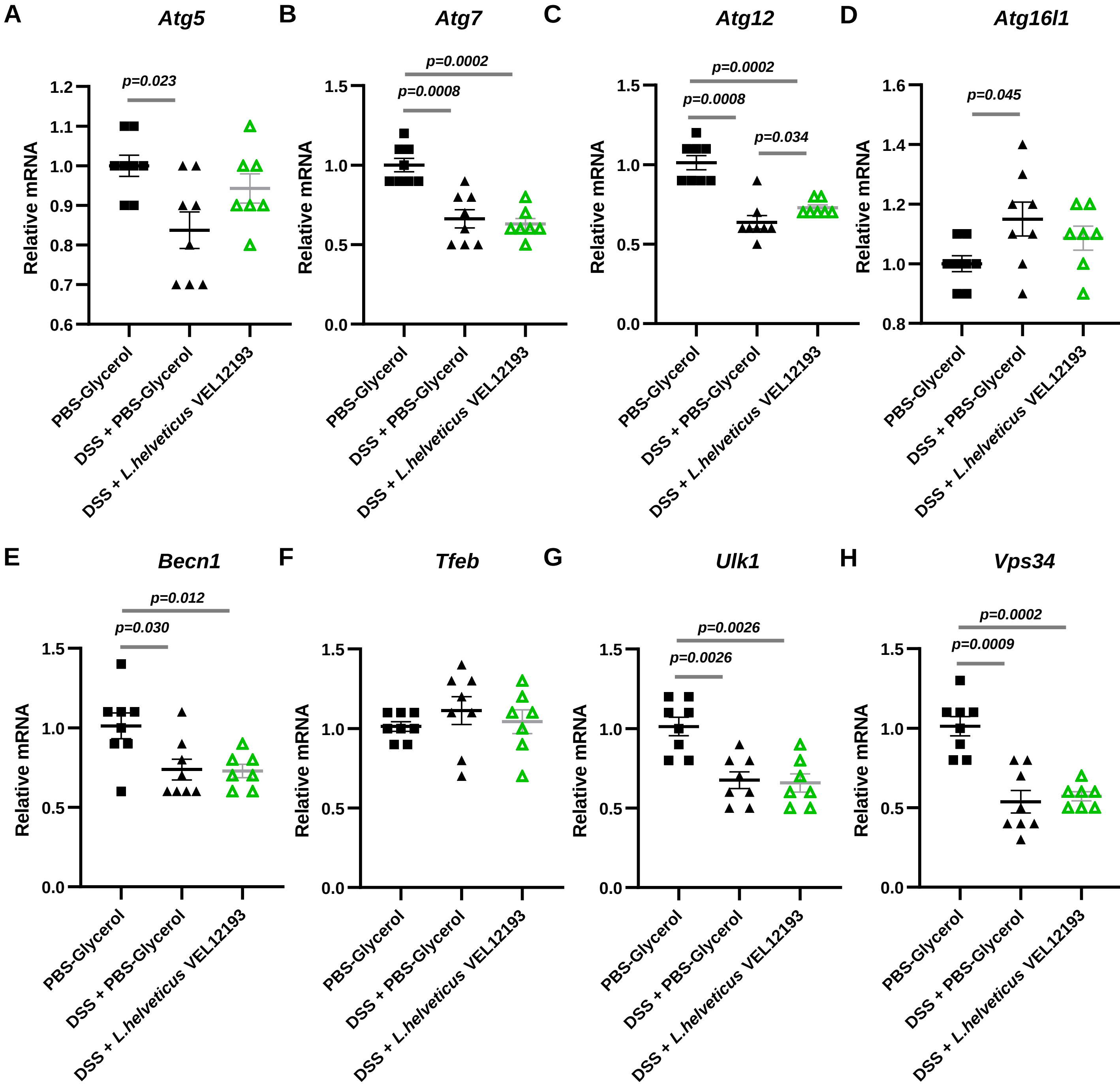


**Supplementary figure 4**
